# A 3D endometrium-on-a-chip reveals the role of conceptus-derived factors CAPG and PDI in conceptus-endometrial communication

**DOI:** 10.1101/2024.09.04.611146

**Authors:** Haidee Tinning, Dapeng Wang, Niamh Forde

## Abstract

Early embryo loss affects all mammalian species, including humans and agriculturally important food-producing mammals such as cattle. The developing conceptus (embryo and extra-embryonic membranes) secretes factors which modify the endometrium and can be critical for early pregnancy processes such maternal recognition of pregnancy (MRP) and enhancing uterine receptivity to implantation. For example, a competent bovine conceptus secretes IFNT to initiate MRP. The bovine conceptus also secretes other proteins at the time of MRP, including CAPG and PDI, which are highly conserved among placental mammals. We have previously shown that these proteins act upon the endometrium to modulate receptivity, embryo development, and implantation in species with different implantation strategies (humans and cattle). We hypothesise that developing a novel 3D bovine endometrium on a chip system will enhance our understanding of the role of conceptus-derived factors in altering the endometrium and/or ULF secretion. Here we have developed a 3D bovine endometrium on a chip system, comprising both stromal and epithelial cell culture combined with culture medium flow better mimics the *in vivo* endometrium and exposure to conceptus-derived factors than conventional 2D endometrial cell culture. We have demonstrated that the conceptus-derived proteins CAPG and PDI modulate the endometrial transcriptome and secretory response to promote pathways associated with early pregnancy and alter ULF composition. This work highlights the critical need for more robust and *in vivo* -like culture systems to study endometrial-conceptus interactions *in vitro* to further investigate the role of conceptus derived factors for pregnancy success.

**SIGNIFICANCE STATEMENT:** We have developed an *in vitro* 3D bovine endometrium-on-a-chip system comprising both primary stromal cells under static conditions and epithelial cells under flow conditions to mimic the *in vivo* endometrial environment from the conceptuses perspective. The secretome of the 3D endometrium-on-a-chip was characterised, was found to contain proteins associated with cell adhesion and tissue development, and contained proteins previously identified in *in vivo* uterine luminal fluid. PDI and CAPG (previously identified conceptus-derived factors) altered the transcriptome and secretome of cells within the system. Exposure to CAPG or PDI altered the secretome of proteins previously identified in pregnant uterine luminal fluid or associated with early pregnancy, and exposure to CAPG or PDI also altered the transcriptome to support processes such as immune response, secretion, proliferation, and adhesion related pathways. This data supports previously published works and highlights the need for the use of more *in vivo-*like *in vitro* models to study conceptus-endometrial interactions.

## INTRODUCTION

The endometrium is the highly heterogeneous specialised tissue lining the internal cavity of the uterus and primarily composed of epithelial and stromal cell types, but also contains microvasculature, immune cells, and stem cells in some species (Cousins et al., 2021). The epithelial cells form a complete monolayer lining the internal cavity of the uterus and can be sub-categorised into luminal or glandular epithelial cells. Stromal cells lie below the epithelial monolayer, making up the bulk of the endometrial tissue (Atkinson et al., 1984).

The epithelial cells, and particularly the glandular epithelial cells (in the form of uterine glands), secrete histotroph into the uterine cavity which contributes to the uterine luminal fluid (ULF) found within the uterine cavity. The ULF can also contain secretions from the oviductal cells (Ghersevich et al., 2015) and from an embryo/conceptus itself if present (Bazer et al., 2011; Forde et al., 2015). The ULF is the source of nutrition for the developing embryo/conceptus prior to placentation (Burton et al., 2002). Studies involving endometrial epithelial gland knockouts in both mouse and sheep have demonstrated the critical importance of endometrial glands supporting early pregnancy-particularly conceptus elongation and the implantation process (Gray et al., 2002; Kelleher et al., 2016; Kelleher et al., 2018).

Much of what is known about early pregnancy in mammals has been achieved using *in vivo* animal studies or traditional *in vitro* techniques which involves the culture of cells (either primary or immortalised). *In vivo* studies require large numbers of animals to be sufficiently powerful to produce statistically significant results, are extremely expensive, often require ethical approval, and require a lot of hands-on work and sample processing (Hartung, 2008). *In vivo* techniques may also not be able to identify low abundant molecules (such as conceptus-secreted factors) due these being diluted by bodily fluids (such as ULF). Although invaluable, *in vivo* animal work can be difficult to achieve and to interpret the results due to variability between animals and between study design (Hartung, 2008). In addition, initiatives such as NC3Rs aim to reduce, replace, and refine the use of animals in research (NC3Rs/BBSRC/Defra/MRC/NERC/Royal Society/WellcomeTrust, 2019), and as such alternatives to *in vivo* animal research should be explored at every opportunity.

As an alternative to *in vivo* models*, in vitro* cell culture studies are well established and routinely used in many laboratories, can be high throughput, and don’t usually require ethical approval. However, it has been demonstrated that cells grown in this manner experience abnormal proliferation and differentiation (Cacciamali et al., 2022). Cell culture growth medium is usually added to a culture vessel and left static for a period (often 48 hours or more), resulting in depleted nutrient availability and increased metabolic waste exposure (Vis et al., 2020), neither of which represents most biological systems. Additionally, conventional cell culture is conducted on a 2D flat culture ware surface, with adherent cells growing in a monolayer (Cacciamali et al., 2022). This doesn’t well recapitulate most biological tissues or systems, including the endometrium. Tissues are more complex-containing multiple cell types in a three-dimensional (3D) conformation which contributes to their function. Innovative approaches to the *in vitro* culture of mammalian cells aim to overcome some of these limitations and bridge the gap between *in vivo* animal studies and *in vitro* traditional culture techniques (Kapałczyńska et al., 2016; Cacciamali et al., 2022).

Advances in 3D modelling approaches have identified and produced *in vitro* models that better recapitulate the *in vivo* tissue structure or environment (Kapałczyńska et al., 2016). Recent examples include microfluidic systems (Young and Beebe, 2010), organ-on-a-chip devices (Leung et al., 2022), scaffolds/extracellular matrix culture supports (MacKintosh et al., 2015), and porous membranes (Chaney et al., 2021). These novel techniques have been developed to overcome the limitations to conventional *in vitro* cell culture and can be applied to investigating endometrial-conceptus interactions. Microfluidic systems provide the opportunity to produce systems where fluid can be pushed through a small chip/channel at a set rate of flow (Young and Beebe, 2010). The channel must be less than 1 mm width or depth to be classified as microfluidics devices (Tiwari et al., 2020). Initially designed to study the physics of fluid movement, microfluidics can be combined with cell culture by loading cells into the chip or channel before applying flow and using culture medium as fluid within the system. Microfluidics can be used to recapitulate a flow rate similar to that of the system being investigated; for this reason microfluidics is often used in the study of blood vessel formation and function under high shear stress (Chu et al., 2023). Culturing cells in medium that is continuously replenished better mimics the *in vivo* environment, as *in vivo* the cells are continuously nutritionally supplied and metabolic waste removed by the circulatory system. By using a device designed to culture multiple cell types, usually in a 3D structure or in series (i.e. different cell types in a series of compartments), it is possible to recapitulate a certain organ or system (Leung et al., 2022). Organ-on-chip (OoC) approaches have been developed for many aspects of reproduction, including the oviduct (Ferraz et al., 2018), placenta (Blundell et al., 2016), and the process of implantation (Park et al., 2022), and was recently reviewed in detail (Young and Huh, 2021). Many groups have fabricated endometrial microfluidic systems in humans, including a 3D system incorporating endometrial epithelial, stromal, and microvasculature cells under flow (Ahn et al., 2021). A more complex example of a endometrium bioengineering is the development of a multi-organ-on-a-chip system which recapitulates the human menstrual cycle through a series of compartments containing ‘on-a-chip’ versions of the ovary, fallopian tube, uterus, cervix and liver (Xiao et al., 2017). The system was shown to develop follicles *in vitro,* secrete steroid hormones, and tissues maintained their *in vivo* like structures *in vitro* (Xiao et al., 2017). Specifically in bovine, a recent study described a system whereby bovine endometrial stromal and epithelial cells could be co-cultured, with stromal cells exposed to varying concentrations of glucose and insulin, to mimic the endometrial exposure to factors in the maternal circulation (De Bem et al., 2021).

We therefore aimed to use a 3D organ-on-a-chip bovine endometrium in combination with microfluidics, to study the conceptus-endometrial communication which occurs in vivo around the critical period of MRP. Specifically, we tested the hypothesis that a 3D cell culture microfluidics approach will allow us to understand the functional roles of conceptus-derived proteins during pregnancy recognition. To test this hypothesis, we aimed to 1) Develop an *in vitro* 3D bovine endometrial model in a microfluidic system comprising both epithelial and stromal endometrial cells, 2) Identify the secretome of the on-a-chip bovine endometrium to compare to *in vivo* ULF and 3) Utilise the endometrium-on-a-chip to investigate how the addition of the conceptus-derived proteins impacts the endometrial transcriptome and secretome.

## MATERIALS & METHODS

Unless otherwise stated all materials were sourced from Sigma-Aldrich.

### Seeding endometrium-on-a-chip devices

Bovine endometrial epithelial (bEECs) and bovine endometrial stromal (bESCs) cells were isolated from uterine tracts obtained from the local abattoir as described in detail in (Tinning et al., 2020). Three uterine tracts in the late-luteal stage of the oestrus cycle were selected based on the morphology of the ovaries to represent the appropriate stage of the cycle where the endometrium is receptive and would be exposed to conceptus-derived factors (Ireland et al., 1980). bESCs and bEECs were cultured in complete bovine medium (RPMI 1640, 10% dextran-coated charcoal-stripped FBS [PAA Cell Culture Company], 1% ABAM) and purified through trypsinisation for 13-14 days to generate epithelial-enriched and stromal-enriched cell populations. Cells were then visually assessed for purity via light microscopy to be over 95% enriched for either bESCs or bEECs. Cells were trypsinised, washed in PBS, and resuspended in complete bovine medium with 10% exosome-depleted FBS (Gibco). bESCs were adjusted to 200,000 cells/mL and bEECs were adjusted to 1,000,000 cells/mL. Fifty-five μL (11,000 cells) of the bESC solution was added the upper static chamber of a μ-Slide Membrane ibiPore Flow ibiTreat microfluidic device (Ibidi, 0.5 μm porous glass membrane, 20% porosity) and incubated for one hour (38.5°C/5% CO_2_) to facilitate cell adherence (bESCs adhered to the top of the membrane). Two hundred µL bEEC solution (200,000 cells) was then added to the lower chamber. All caps were added to the inlets/outlets to prevent evaporation, immediately inverted (so bEECs adhered to the underside of the membrane), placed into a sterile petri dish, and incubated (38.5°C/5% CO_2_) overnight. The microfluidic device was then de-inverted, the lower chamber caps and medium removed, and replenished with exosome-depleted bovine medium (detailed above). The chip was then returned to the incubator (38.5°C/5% CO_2_) for 3-4 days until bEEC layer was 90% confluent. Medium was replenished every two days from the lower channel inlet.

### Microfluidic flow treatment system

Recombinant bovine forms of CAPG (rbCAPG) and PDI (rbPDI) were produced as described (Tinning et al., 2020; Tinning et al., 2024) by Newcastle Universities Protein and Proteome Analysis Facility (UK) and purified into PBS. Treatments were prepared in exosome-depleted complete bovine medium as follows: 1) Vehicle control (VC-PBS), 2) rbCAPG 1000 ng/mL, or 3) rbPDI 1000 ng/mL. Concentrations were previously shown to alter the transcriptome in 2D static culture systems (Tinning et al., 2020; Tinning et al., 2024). Treatments were loaded into 5 mL leur lock syringes (Terumo), capped, and pre-warmed in the incubator to 38.5°C overnight. Bubbles were then removed from the syringes by tapping. A shelf was removed from the incubator, cleaned with 70% ethanol, and placed into a sterile laminar flow hood. A syringe pump (New Era Pump Systems) was wiped with 70% ethanol and placed into the hood on the incubator shelf. The pre-warmed syringes were secured in the pump system securely and pre-prepared 0.8 mm ID sterile silicone tubing (Ibidi) was attached with a luer lock female connector (Ibidi) to the syringe. The tubing was then attached to an elbow luer connector (Ibidi) and the syringes manually pushed to fill the tubing with medium until a droplet appeared at the outlet of the elbow leur connector. The devices were removed from the incubator and the lower channel flushed with PBS three times by adding to the inlet and removing from the outlet. The lower channel inlet and outlet was then overfilled to produce a dome of liquid at the surface. The droplet leaving the tubing from the syringe was then connected to the domed liquid inside the lower channel. This linking process was necessary to prevent bubbles. The elbow luer connector male was then pushed on a 90-degree angle into the device lower channel inlet and turned back 90 degrees to seal. A short piece of sterile tubing was pre-prepared with a second elbow luer connector male attached to one end. To the other end a 1 mL syringe (BD Plastipak) with a blunt needle (SOL-Millennium) was inserted and used to push PBS into the tubing until a droplet formed from the outlet of the elbow luer connector male. This droplet was then linked to that of the devices domed medium of the lower channel outlet, again to prevent bubbles being trapped inside the microfluidic system. The needle was simultaneously removed from the outlet tubing whilst pushing the outlet elbow luer connector male into the chip outlet at a 90-degree angle and sealing by twisting back 90 degrees. The syringes were then very gently pushed slowly to inject medium through the chip and microfluidic tubing system until the whole system was filled with medium. The end of the outlet tubing was then pushed into a hole made in the lid of a seven mL sterile bijou to collect the conditioned medium during flow. The complete flow system on the shelf was transferred back to the incubator (38.5°C/5% CO_2_). The pump device was then plugged in, and flow rate set to 0.8 μL/min and set to run for 24 hours to mimic what is produced in vivo, as described below. Two one mL samples of VC medium were also placed in the incubator alongside the system for 24 hours in duplicate (unconditioned medium). A graphical representation of the 3D endometrium on a chip is seen in Figure 1.

**Figure 1.**
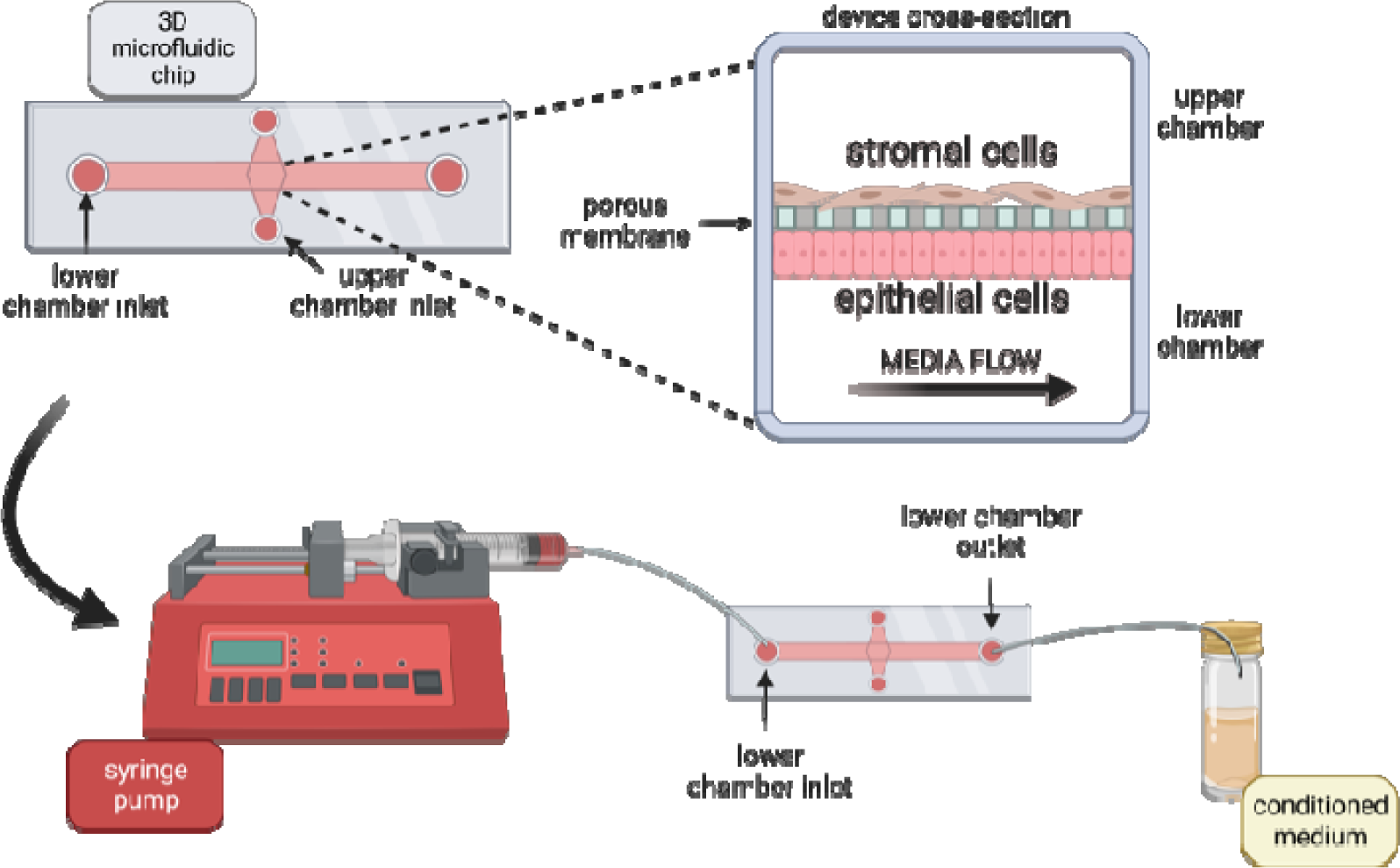
Design of 3D bovine endometrial Ibidi organ-on-a-chip microfluidic system. Lower chamber has an inlet on the left, and outlet on the right, which are connected via adaptors and tubing to a syringe containing medium. The syringe is connected to a syringe pump which pushes the syringe plunger to push medium through the lower chamber at 0.8 μL per min. Conditioned medium is collected from the lower chamber outlet. Upper chamber is separated by a porous glass membrane and is static (not under flow). bEECs seeded in lower chamber to underside of membrane and bESCs seeded in upper chamber to confluency. Figure created using biorender.com.

The flow rate of 0.8 μL/min was determined for the 3D endometrium microfluidic channel based on previous experiments ((De Bem et al., 2021; Tinning et al., 2024). An initial 1 μL/min flow rate was chosen because uterine tubal secretions were found to be 1.43mL/24 hours during dioestrus and 1.54mL/24 hours during oestrus in cattle (Segura-Aguilar and Reyley, 2005), with the rate of 1 μL/min equating to 1.44 mL/24 hours. To attempt to match the shear stress experienced by the epithelial cells under flow, the flow rate of 1 μL/min was changed to 0.8 μL/min for the 3D culture system based on the dimensions of the channel.

### Sample collection

After 24 hours the flow of medium was stopped, the pump unplugged, and the whole system moved back to the laminar flow hood. First, the conditioned medium (and unconditioned samples) were transferred to 2 mL sterile microfuge tubes and stored at 4°C until processing. Next, the syringes were detached from the tubing and all connectors were uncoupled from the devices and the tubing separated. The devices were flushed with PBS, the lower flow channel was flushed three times. The upper channel was flushed with 55 μL PBS by using two P200 pipettes, one to push in 55 μL PBS to the inlet and the other to simultaneously extract 55 μL liquid from the outlet-this was also repeated three times. At this point the devices were visually inspected under a light microscope to ensure the membrane was intact before continuing. Both the upper and lower channels were filled with tryspin solution (0.025% for bESCs or 0.25% for bEECs) and incubated for 3 mins. Exosome-depleted complete bovine medium was added to each channel and cells collected from the outlets. Device was checked via light microscopy for any remaining adhered cells and to ensure the membrane was still intact, and the trypsinisation processes repeated if needed. Collected cells were centrifuged to pellet at 500 g for 5 mins, resuspended in 1 mL PBS to wash, and re-pelleted at 500 g for 5 mins. Supernatant was aspirated gently and cell pellets snap-frozen in liquid nitrogen and transferred to a −80°C freezer.

The conditioned medium collected from the microfluidic lower-chamber flow-through, and the unconditioned medium samples, were processed to remove debris. Firstly, the medium was centrifuged at 500 g for 10 mins to pellet cells. Supernatant was carefully transferred to a new microfuge tube and the pellet discarded. Next, the supernatant was subjected to a 2000 g 10 min spin to pellet cell debris. Supernatant was carefully transferred to a new Eppendorf and the pellet discarded. Finally, the supernatant was centrifuged at 14,000 g for 30 mins to pellet microvesicles. The resulting supernatant was snap-frozen and stored at −80°C.

### Transcriptome analysis

RNA was extracted from both the bEEC and bESC pellets using the miRNeasy Micro Kit (Qiagen) as per manufacturers’ instructions, snap-frozen in liquid nitrogen, and stored at −80°C.RNA libraries were prepared and sequenced by Novogene (Cambridge) as per their standard protocols which are described here based on information provided. Due to the small amount of RNA recovered, a SMARTer amplification process was performed using SMART-Seq v4 Ultra Low Input RNA Lit for sequencing (Clontech) to synthesise the double-stranded cDNA libraries. Briefly, first strand cDNA synthesis was performed, followed by template switching and extension, then cDNA amplified by PCR to produce double stranded cDNA. The amplified cDNA samples were purified with AMPure XP beads and quantified with a Qubit 2.0 fluorometer (Life Technologies). Library prep was then carried out using the Novogene NGS RNA library prep set (PT042). Briefly, the cDNA samples were sheared by the Covaris system, then sheared fragments underwent end-rapir, A-tailed, and ligated to sequencing adaptors. During this process a 200 bp size selection is used. The resulting libraries was then checked with Qubit 2.0 fluorometer (Life Technologies), diluted to 2 ng/μL, insert size checked on an Agilent 2100 bioanalyser and quantified to greater accuracy by qPCR. Quantified libraries were then pooled and sequenced using the Illumina NovaSeq 6000 machine (Illumina, California, USA) with a paired end 150bp length read. RNA sequencing raw data was processed and differentially expressed genes (DEGs) as described in detail previously (Tinning et al., 2020). Briefly, reads were aligned to the reference genome and gene annotation files of cow (*Bos taurus*) from Ensembl genome database (release 96) (Cunningham et al., 2019). Reads were mapped using Rsubread (Yang Liao et al., 2019), and reads quantified using featureCounts. Statistical test for differential gene expression was conducted via DESeq2 (Love et al., 2014) with the cut-offs such as log2FoldChange >1 (or <-1) and padj <0.05. For PCA plotting of each group of samples, protein-coding genes with RPKM value ≥ 1 in at least one sample were used and subsequently log2(RPKM+1) transformation and a quantile normalisation were applied. Only protein-coding genes and lncRNAs were retained for further analysis. Overrepresentation enrichment analysis of differentially expressed protein-coding gene sets was executed using WebGestalt (webgestalt.org) (Yuxing Liao et al., 2019). For enriched gene ontology terms, biological process non-redundant datasets were chosen as the functional database, *Bos taurus* selected as species of interest, and significance level was determined by FDR <0.05. Venn diagram analysis was performed using Venny 2.1.0 (bioinfogp.cnb.csic.es) (Oliveros, 2007).

### Proteomics analysis of conditioned medium

The processed conditioned medium was depleted of bovine albumin using an albumin depletion kit, according to the manufacturer’s protocol (Thermo Fisher Scientific, UK). An equal volume of each depleted sample (equivalent to 20-50µg protein) was then digested with trypsin (1.25 µg trypsin; 37°C, overnight), labelled with Tandem Mass Tag (TMT) eleven plex reagents according to the manufacturer’s protocol (Thermo Fisher Scientific, UK) and the labelled samples pooled. The pooled sample was then processed and underwent nano-LC mass spectrometry as described previously described (De Bem et al., 2021). The raw data files were processed and quantified using Proteome Discoverer software v2.1 (Thermo Scientific) and searched against the UniProt Bos taurus database (downloaded September 2020: 46224 entries) using the SEQUEST HT algorithm. Peptide precursor mass tolerance was set at 10ppm, and MS/MS tolerance was set at 0.6Da. Search criteria included oxidation of methionine (+15.995Da), acetylation of the protein N-terminus (+42.011Da) and Methionine loss plus acetylation of the protein N-terminus (−89.03Da) as variable modifications and carbamidomethylation of cysteine (+57.021Da) and the addition of the TMT mass tag (+229.163Da) to peptide N-termini and lysine as fixed modifications. Searches were performed with full tryptic digestion and a maximum of 2 missed cleavages were allowed. The reverse database search option was enabled, and all data was filtered to satisfy false discovery rate (FDR) of 5%. The resulting list of proteins for each sample were then analysed in Excel to determine the fold change in protein abundance between treatment groups and the associated p-value (Aguilan et al., 2020). Fold changes were calculated by taking the average of the vehicle control samples from the average of the rbPDI/rbCAPG treated conditioned medium samples. Fold changes for the ‘*in vitro* ULF’ were calculated by taking the average of the unconditioned medium samples from the average of the vehicle control samples. Proteins with a positive fold change are therefore more abundant and those with a negative fold change value are less abundant. For rbCAPG/rbPDI samples, P-value significance was calculated by paired two-tailed t-test. All comparisons with a p-value <0.05 was considered significantly different. For the *in vitro* ULF samples, first an F-test was carried out to determine which proteins displayed a different variance between conditioned and unconditioned samples, as the samples were not paired. For the proteins which had an F-test value p<0.05 a two-tailed two-sample unequal variance t-test was performed, and those that had an F-test value p>0.05 a two-tailed two-sample equal variance t-test was performed. Full data shown in Supplementary Table S1. Protein lists were then filtered for p<0.05 and excluded any that were identified as false for bovine and/or true for contaminant.

### Proteomics data downstream analysis

To firstly investigate the data, the list of proteins and imputation values (post-normalisation, shown in Supplementary Table S1, were placed into a .txt table and loaded into RStudio version 4.2.2 (Team, 2021). A box plot was produced using BioStatR version 4.0.1 (Bertrand and Maumy-Bertrand, 2023). A PCA plot was produced using ggfortify version 0.4.16 (Tang et al., 2016). Next, to visualise the *in vitro* ULF data, a volcano plot was produced using the EnhancedVolcano Bioconductor package version 1.16.0 (Blighe K, Rana S, 2023). This list of *in vitro* produced ULF proteins identified were then compared to those identified *in vivo* in other studies, including Forde *et al* (2015). Data by Forde *et al* was taken from their Supplemental table S1, which identified 334 proteins present in the ULF from at least 3 out of 4 day 16 non-pregnant cattle. The list of 334 proteins were identified by a ‘NCBI GI number’, and so were converted to UniProtKB Swiss Prot identifiers to match the identifiers used in this study. One hundred and forty-three proteins were converted to bovine UniProt KB Swiss Prot identifiers using the UniProt ID mapping online tool (uniprot.org/id-mapping) (Huang et al., 2011). Enrichment and protein-protein association analysis was carried out using STRING DB (string-db.org) (Szklarczyk et al., 2019).

## RESULTS

### 3D bovine endometrium-on-a-chip secretome

Following proteomics analysis of the conditioned medium, principal component analysis (PCA) was used to visualise the spread of the data after normalisation. The PCA plot (Figure 2A) showed the conditioned medium samples cluster mostly by biological replicate (in PC1), with the two unconditioned medium samples clustered together at the top of the plot (in PC2). The data was then investigated to assess the spread of the individual data points using a box plot (Figure 2B) demonstrating the data was similar across all sample types. To determine what *in vitro* ULF secretion occurred, proteins present in the conditioned vehicle control samples produced from the 3D endometrium-on-a-chip microfluidics system that were significantly increased or decreased (p<0.05) in abundance when compared to the unconditioned medium samples were identified (Figure 3A; Supplementary Table S2). Those that were significantly increased in abundance were then identified (Table 1) and were collectively termed ‘*in vitro* ULF’ as they were secreted into the conditioned medium in this endometrial model system. Of the 68 enriched *in vitro* ULF proteins, 61 were mapped by STRING DB. STRING DB interaction analysis of the 61 proteins demonstrated that many of the proteins secreted from the 3D endometrial chip are associated and/or interact with each other (i.e. have edges connecting them to other proteins) (Figure 3B). There was one large cluster and a small unconnected cluster of four proteins (names taken from STRING DB) (Signal peptidase complex subunit 2 [SPCS2], ribosome binding protein 1 [RRPB1], Atlastin GTPase 3 [ATL3], and Reticulon [RTN4]).

**Figure 2.**
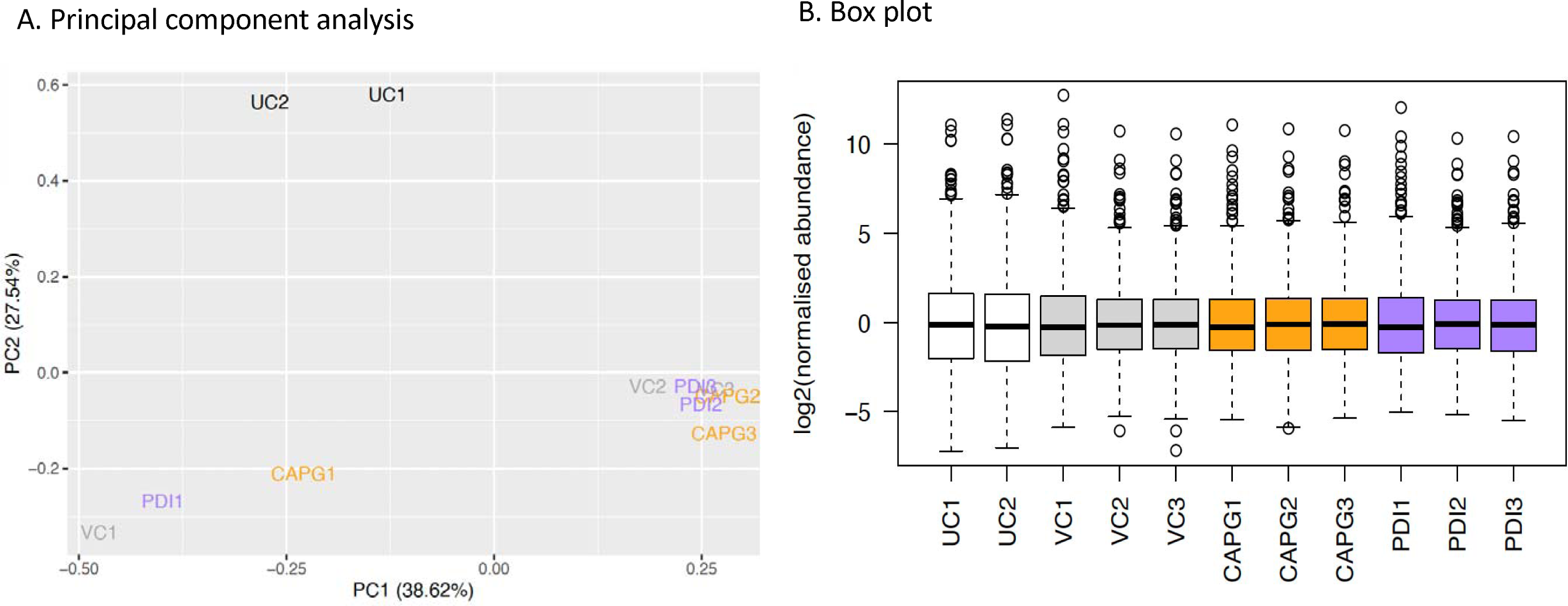
Overall proteomic analysis of conditioned medium in 3D endometrium-on-a-chip. A. Principal component analysis plot of conditioned medium from 3D microfluidics device (in vitro ULF). B. Boxplot of normalised abundance values for proteins identified in the in vitro ULF. Conditioned medium flowed through bEEC culture chamber connected via porous membrane to static bESC culture chamber. Treatments were added to culture medium flowed through the bEEC chamber at a rate of 0.8 μL/min for 24 hours and contained one of the following treatments in biological triplicate: UC = unconditioned, VC = vehicle control, PDI = rbPDI 1 μg/mL, CAPG = rbCAPG 1 μg/mL. Numbers following VC, PDI, or CAPG samples indicate biological replicate. Proteins identified in medium using TMT mass spectrophotometry. PCA produced using ggplot in RStudio.

**Figure 3.**
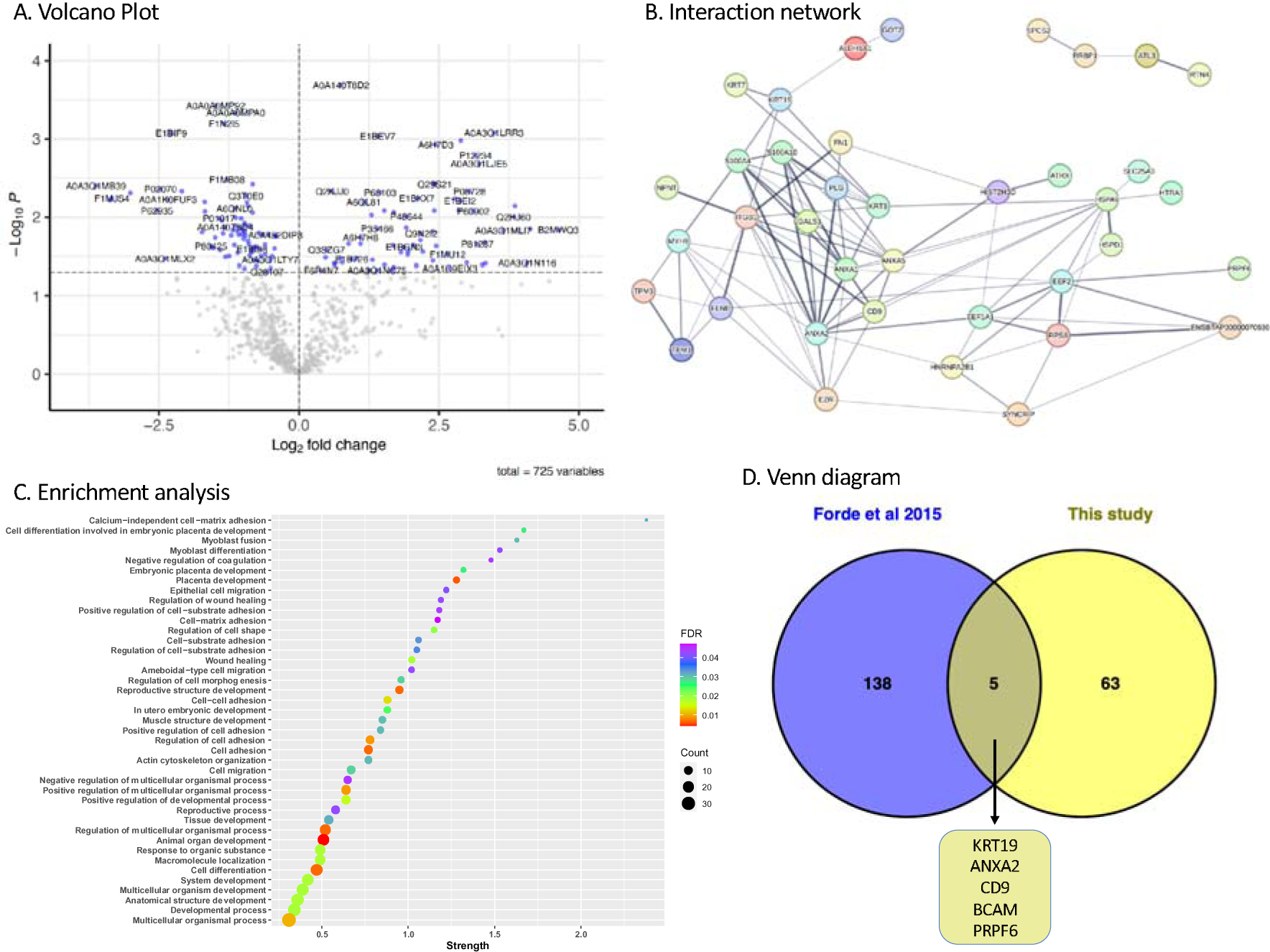
Protein composition of in vitro endometrium on a chip produced ULF. A. Volcano plot of differentially abundant proteins present in conditioned medium (vehicle control) samples compared to the unconditioned medium samples. Blue samples above the dashed line are significantly changed in abundance between the conditioned (n=3) vs unconditioned samples (n=2) (p<0.05). Conditioned medium flowed through bEEC culture chamber at a rate of 0.8 μL/min for 24 hours separated by a porous membrane to a static bESC culture chamber. Figure produced in BioConductor using EnhancedVolcano in R Studio. B. String DB analysis of in vitro ULF proteins. Each node represents a protein, edges (connections) represent functional/physical protein associations with a minimum required interaction score of ‘medium 0.4’. Thickness of edge represents the strength of supporting data. In vitro secreted proteins determined by comparing conditioned medium produced by the endometrium-on-a-chip system (n=3 vehicle controls) flowed through the bEEC chamber connected to a static bESC chamber via a porous glass membrane, to the unconditioned medium (n=2) samples (p<0.05). Produced using STRING DB. Disconnected nodes removed for clarity C. Enriched go terms associated with proteins secreted in the in vitro ULF. Proteins significantly enriched (positive fold change abundance) in conditioned vehicle control medium samples (n=3) compared to unconditioned samples (n=2) (p<0.05), from endometrium-on-a-chip culture system, were subjected to go term enrichment analysis in STRING DB (FDR <0.05). Strength of enrichment is Log10(observed/expected). Graph produced using ggplot in R Studio. Full data in Supplementary table S3. D. Venn diagram comparing in vivo ULF proteins to in vitro secreted ULF proteins. In vivo ULF proteins identified by Forde et al (2015) as present in day 16 non-pregnant cattle compared to in vitro ULF proteins identified as secreted by the 3D endometrium-on-a-chip microfluidic system described here. ULF = uterine luminal fluid. Full data in supplementary table S4.

**Table 1.**
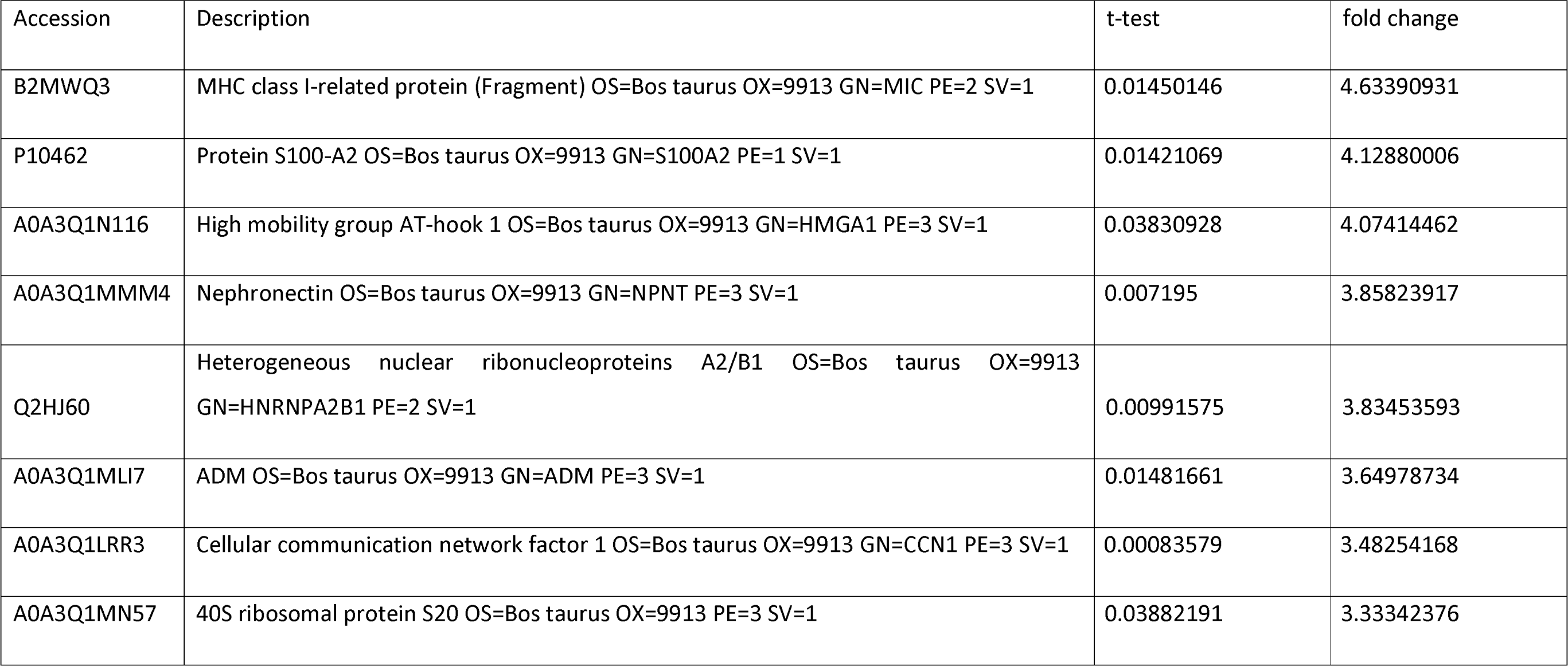
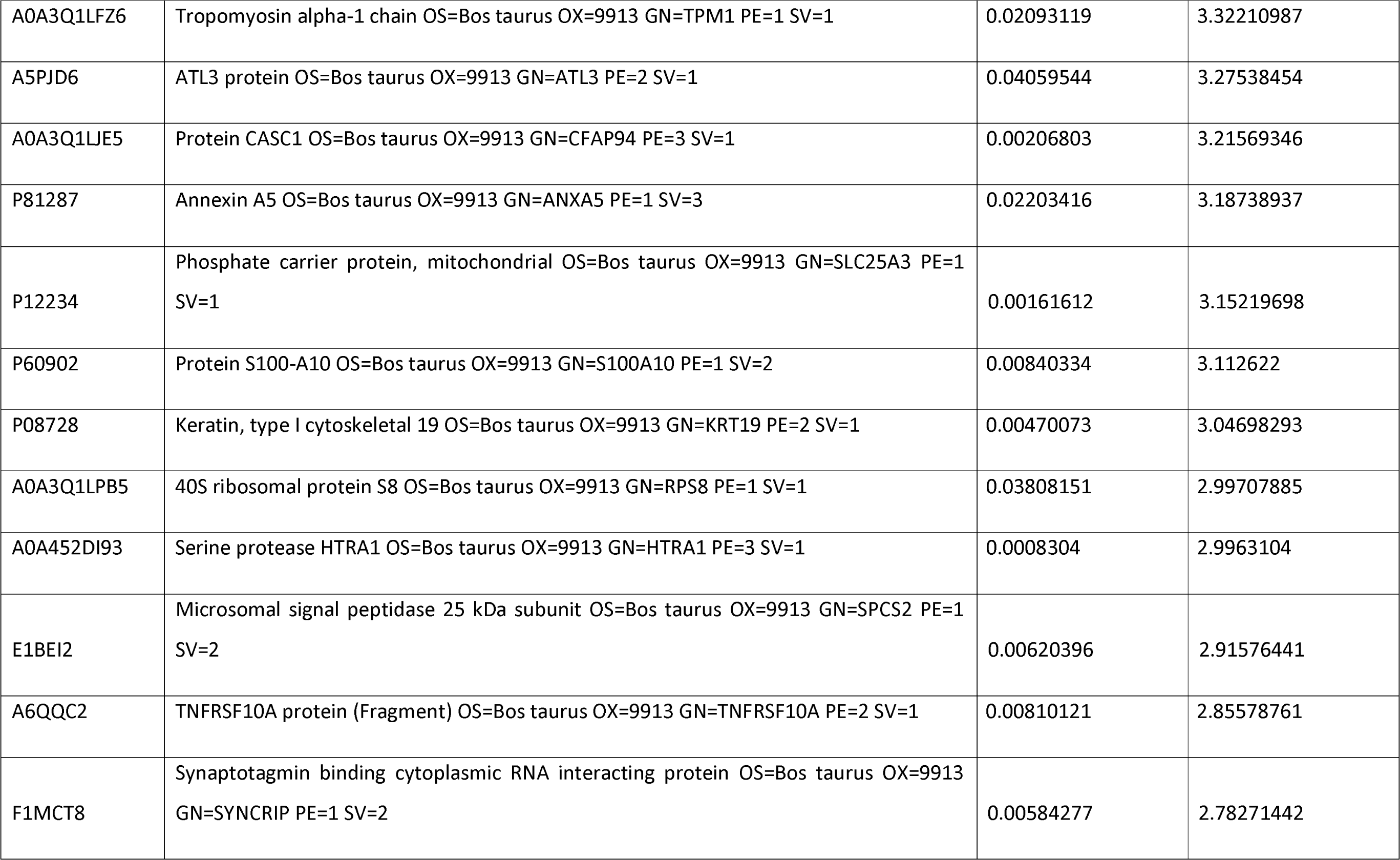
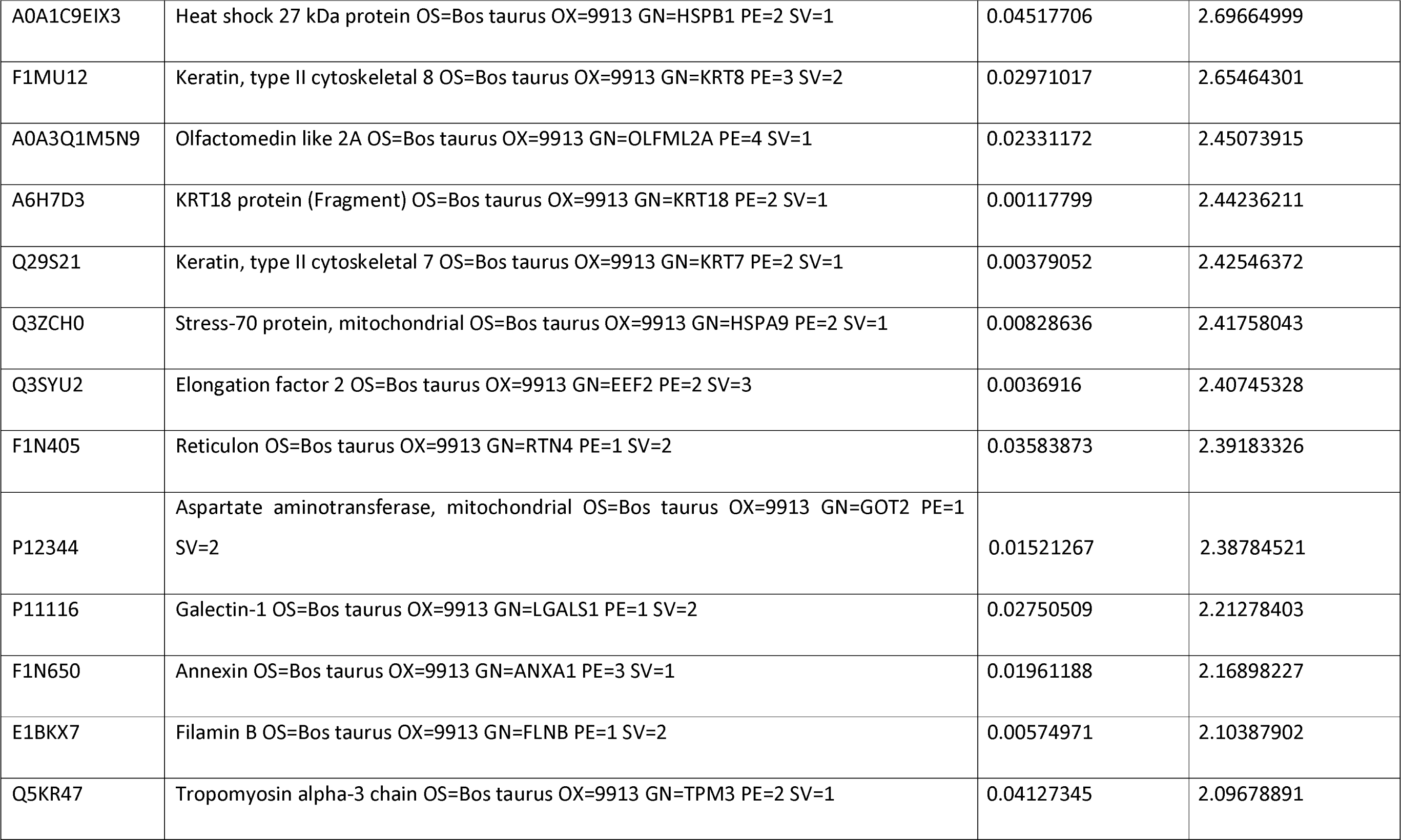
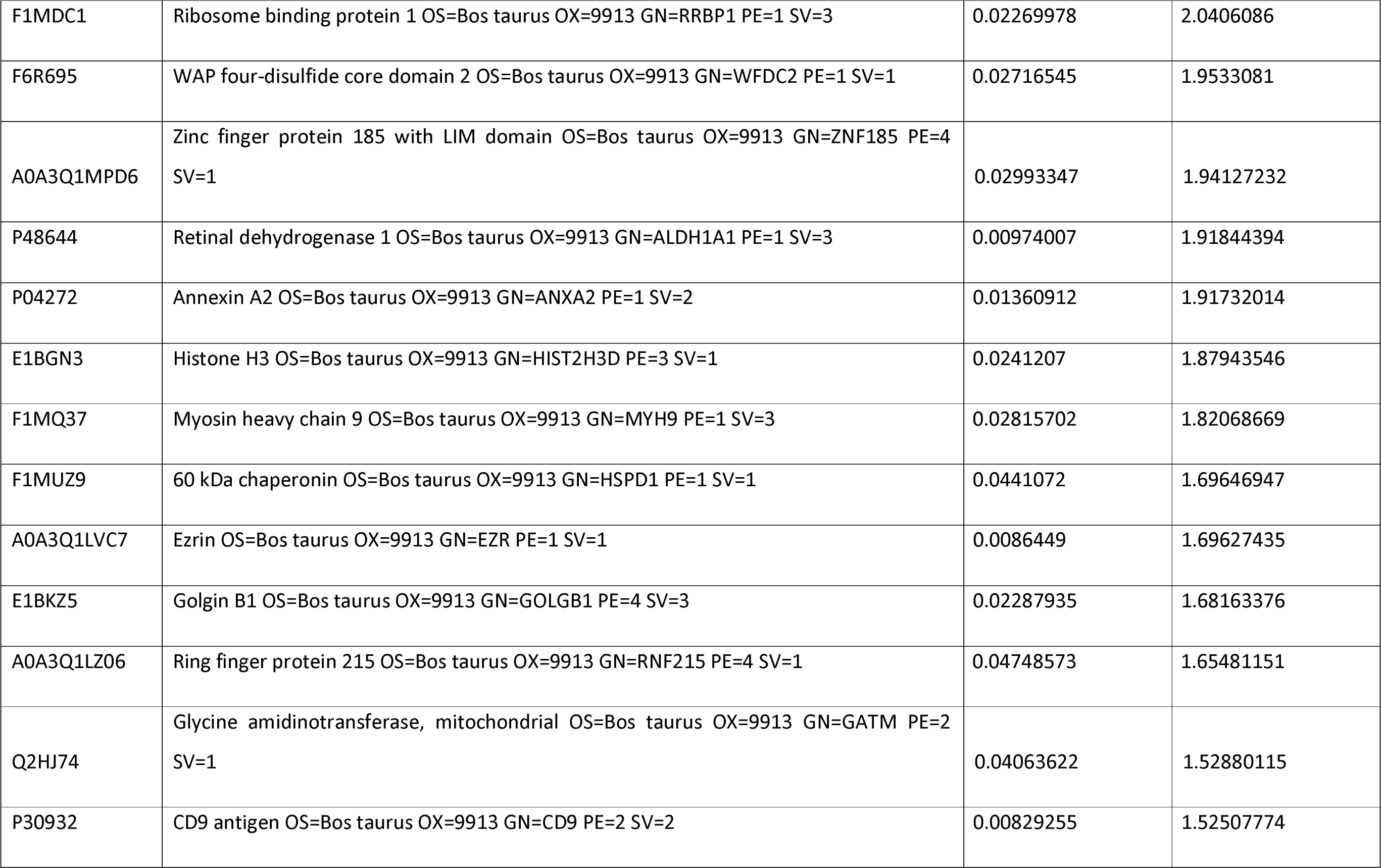
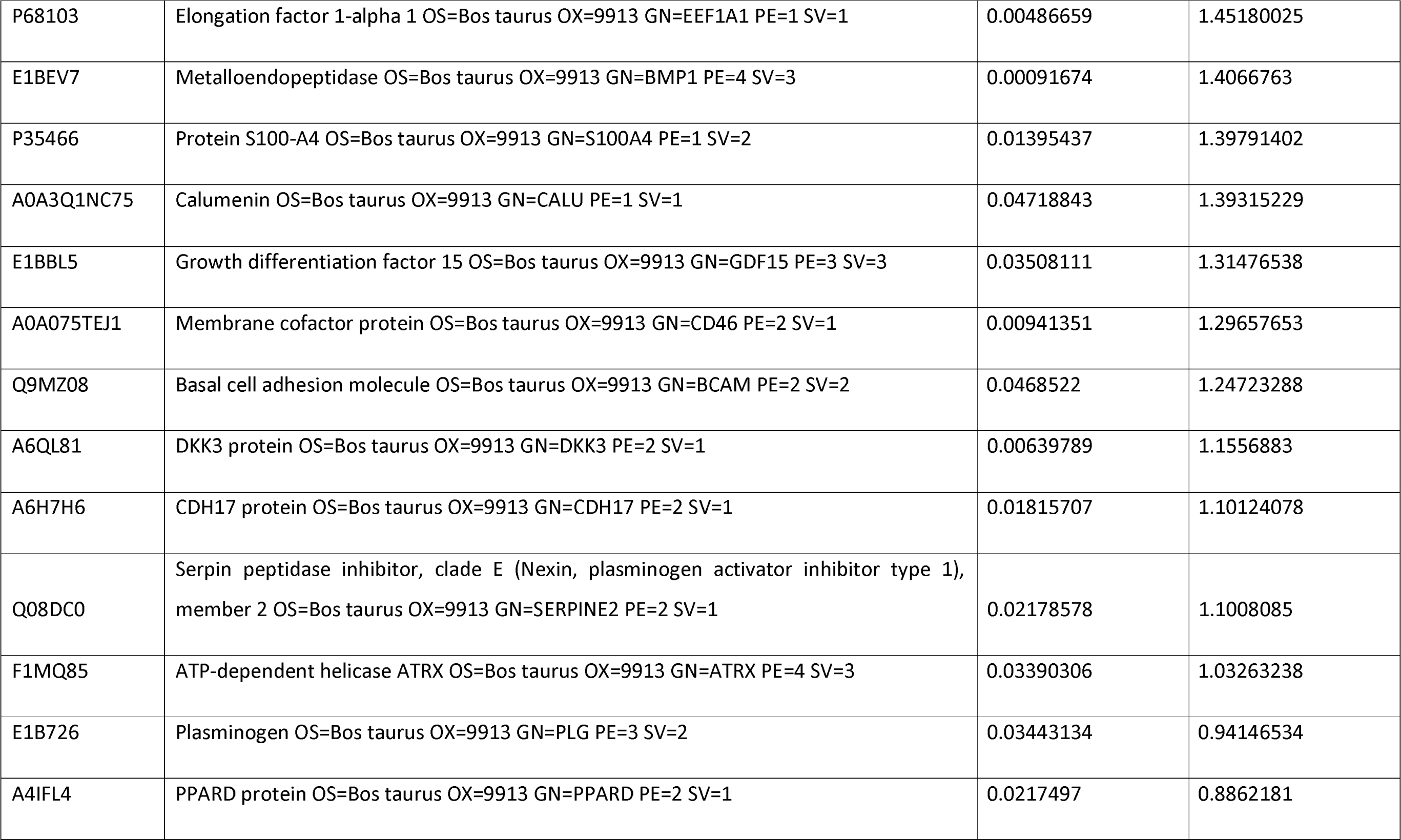
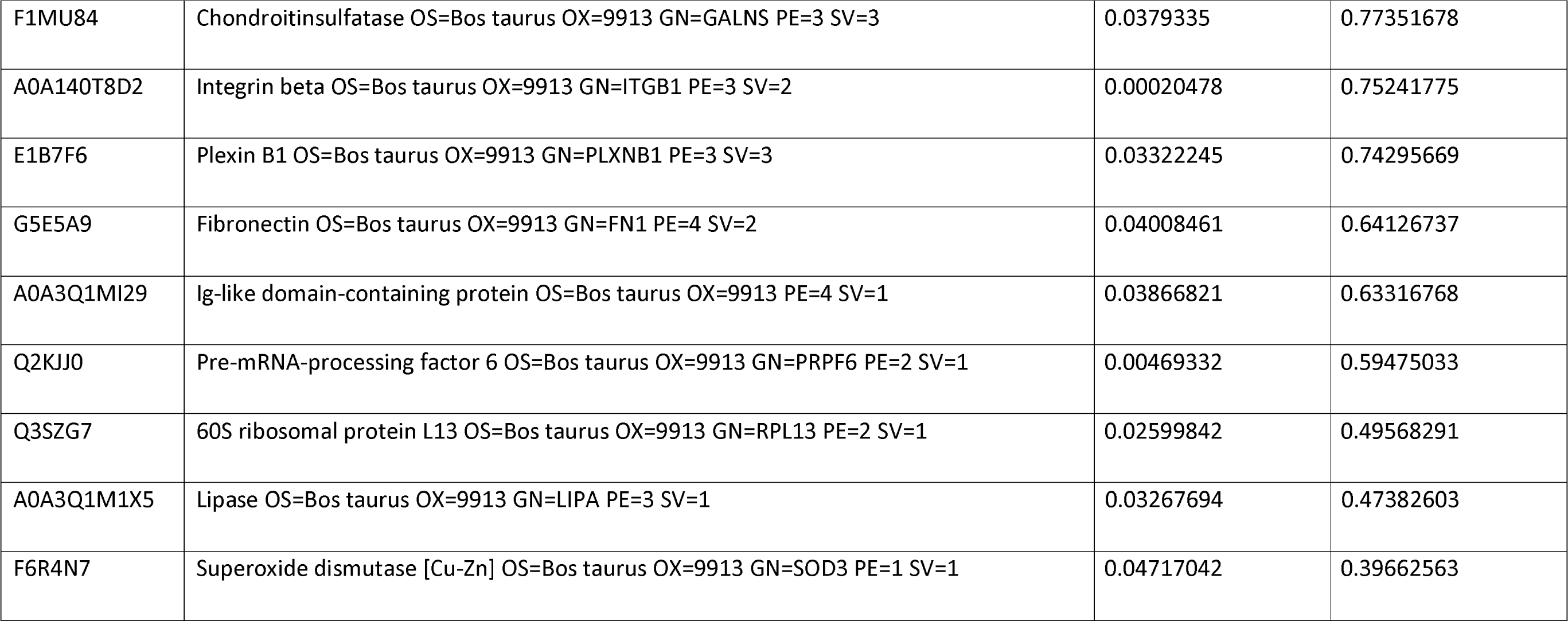
List of ‘*in* vitro’ ULF secreted proteins. Conditioned medium produced by the endometrium-on-a-chip system conditioned medium flowed through EEC chamber. bEEC chamber connected to a static bESC chamber via a porous glass membrane. Enriched proteins present in conditioned medium vehicle control) samples compared to the unconditioned medium (n=2) samples (p<0.05). Fold-change and t-test carried out in Excel as described by an *et al* (2020)

Enrichment analysis revealed biological process go terms associated with proteins secreted in the *in vitro* ULF include calcium-dependant cell-matrix adhesion, cell differentiation involved in embryonic placenta development, and myoblast fusion as the most significantly enriched (Figure 3C: Supplementary Table S3).

To compare the *in vitro* ULF proteins secreted in the 3D endometrium-on-a-chip system to those produced *in vivo,* these data were compared to a key study which used mass spectrophotometry to identify proteins present in *in vivo* non-pregnant bovine ULF on day 16 (Forde et al., 2015). This comparison determined that only 5 of the 68 proteins identified here were also found in *in vivo* ULF on day 16-keratin 19 (KRT19), annexin A2 (ANXA2), CD9, basal cell adhesion molecule (BCAM), and pre-mRNA-procession factor (PRPF6) (Figure 3D: Supplementary Table S4).

### CAPG alters the ability of endometrium to support early pregnancy

The addition of rbCAPG to the microfluidic flow altered abundance of 25 proteins in the conditioned medium compared to vehicle control samples, including the CAPG protein as the most highly abundant (Table 2). STRING DB analysis of the 25 proteins revealed that many nodes had no connecting edges, and therefore no known direct interactions, but revealed some clear main interacting clusters, including Osteoglycin/Mimecan (OGN), Thrombospondin-4 (THBS4), Collagen type VI alpha 1 chain (COL6A1), and Heterogeneous nuclear ribonucleoprotein R (HNRNPR), with Glyceraldehyde-3-phosphate dehydrogenase (GAPDH) seemingly acting as a central connecting proteins known as ‘hub proteins’ (Figure 4A).

**Figure 4.**
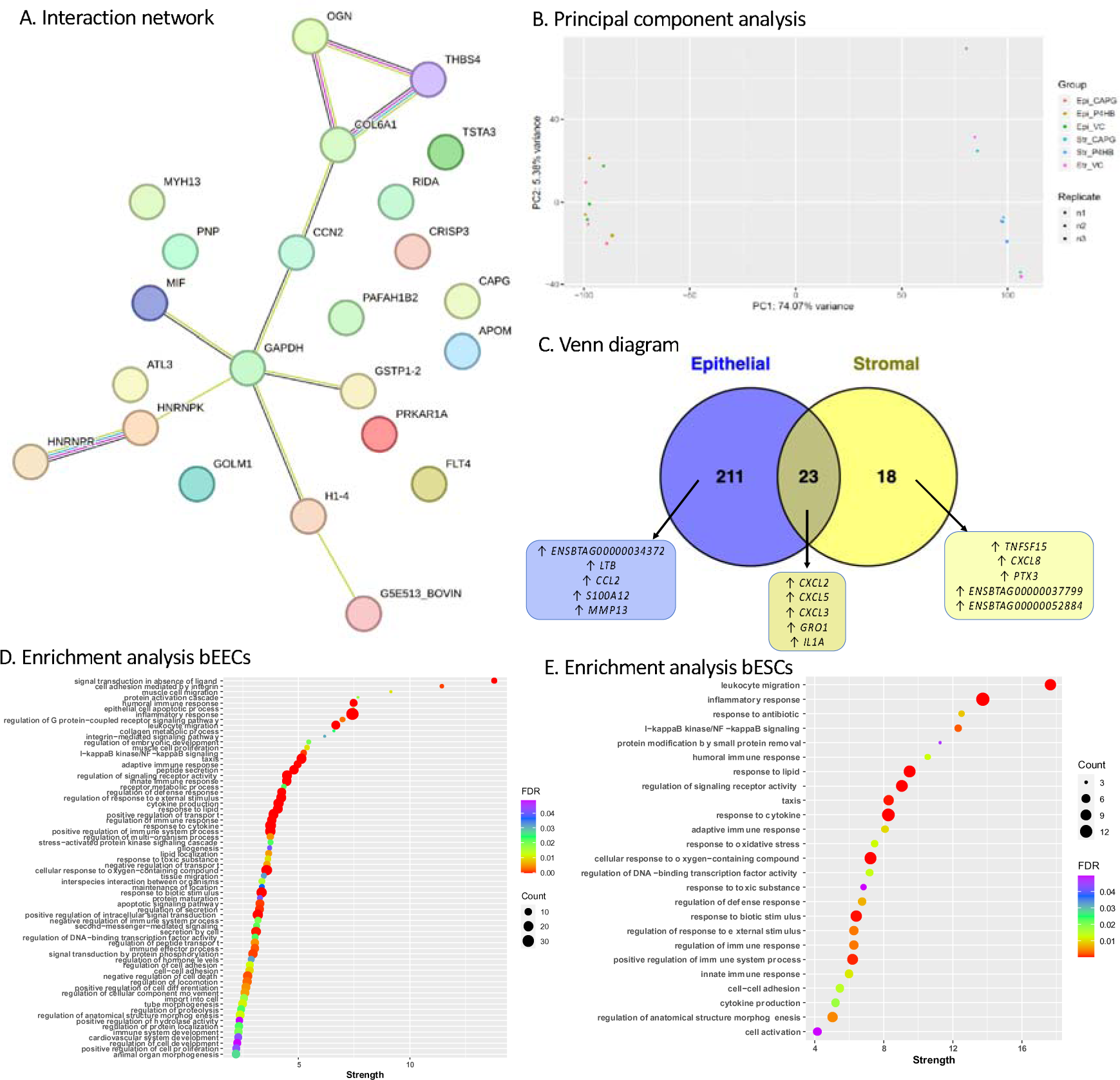
Impact of CAPG on endometrial secretome and transcriptome. A. Interaction analysis of differentially abundant proteins in conditioned medium supplemented with rbCAPG compared to vehicle control samples (p<0.05). Each node represents a protein, nodes with edges represent functional/physical protein associations with a minimum required interaction score of ‘medium 0.4’. CAPG was added to the culture medium in bEEC culture chamber. Thickness of edge represents the strength of supporting data. Produced using STRING DB. B. Principal component analysis of transcriptome of 3D endometrium-on-a-chip epithelial and stromal cells. bEECs and bESCs were cultured in a 3D microfluidic chip, with the bEECs under flow (0.8 μL/min) with culture medium containing rbCAPG (CAPG) or rbPDI (P4HB) 1 μg/mL, or PBS vehicle controls (VC) for 24 hours. Epi_ = epithelial cells, Str_ = stromal cells, VC = vehicle control. C. Venn diagram of differentially expressed genes in response to rbCAPG treatment. rbCAPG treatment in microfluidic flow through resulted in differentially expressed genes in bovine epithelial and stromal cells when compared to vehicle control samples (n=3, padj<0.05, fold change <1 or >1). Produced in Venny. ↑ upregulated, ↓ downregulated. Top 5 up/down regulated genes shown. Full data in Supplementary Table S7. D. Biological processes gene ontologies enriched in bovine epithelial cells inside 3D microfluidic organ-on-a-chip device in response to rbCAPG treatment. Analysis carried out in Webgestalt, FDR<0.05, strength indicates enrichment ratio. Full data in Supplementary table S8. Figure produced using ggplot in R Studio. E. Biological processes gene ontologies enriched in bovine stromal cells inside 3D microfluidic organ-on-a-chip device in response to rbCAPG treatment. Analysis carried out in Webgestalt, FDR<0.05, strength indicates enrichment ratio. Full data in Supplementary table S9. Figure produced using ggplot in R Studio.

**Table 2.** List of differentially abundant proteins in culture medium following CAPG treatment. rbCAPG added to the culture medium flowing through the fluidic bovine endometrium-on-a-chip device for 24 hours and proteins present in the conditioned medium determined by mass spectrophotometry, entially abundant proteins identified compared to vehicle control samples (n=3 biological replicates.

| Accession | Description | t-test | fold change |
| --- | --- | --- | --- |
| A0A3Q1N6D1 | Macrophage-capping protein OS=Bos taurus OX=9913 GN=CAPG PE=1 SV=1 | 0.0060 | 3.8010 |
| Q3T0D0 | Heterogeneous nuclear ribonucleoprotein K OS=Bos taurus OX=9913 GN=HNRNPK PE=2 SV=1 | 0.0170 | 0.8219 |
| A0A452DJA8 | Inosine phosphorylase OS=Bos taurus OX=9913 GN=PNP PE=3 SV=1 | 0.0410 | 0.8061 |
| P10096 | Glyceraldehyde-3-phosphate dehydrogenase OS=Bos taurus OX=9913 GN=GAPDH PE=1 SV=4 | 0.0261 | 0.5961 |
| A5PJD6 | ATL3 protein OS=Bos taurus OX=9913 GN=ATL3 PE=2 SV=1 | 0.0207 | 0.5689 |
| A7E336 | 60S acidic ribosomal protein P0 OS=Bos taurus OX=9913 GN=RPLP0 PE=2 SV=1 | 0.0004 | 0.5343 |
| A0A3Q1NB36 | Heterogeneous nuclear ribonucleoprotein R OS=Bos taurus OX=9913 GN=HNRNPR PE=1 SV=1 | 0.0369 | 0.4246 |
| A0A3Q1M2A1 | GDP-4-keto-6-deoxy-D-mannose-3,5-epimerase-4-reductase OS=Bos taurus OX=9913 GN=TSTA3 PE=1 SV=1 | 0.0151 | 0.3875 |
| G3MWV5 | H15 domain-containing protein OS=Bos taurus OX=9913 GN=H1-4 PE=3 SV=2 | 0.0183 | 0.3157 |
| A0A3Q1LMV5 | GST class-pi OS=Bos taurus OX=9913 GN=GSTP1 PE=3 SV=1 | 0.0267 | 0.2374 |
| E1BI98 | Collagen type VI alpha 1 chain OS=Bos taurus OX=9913 GN=COL6A1 PE=1 SV=1 | 0.0418 | -0.1099 |
| E1BLA8 | Golgi membrane protein 1 OS=Bos taurus OX=9913 GN=GOLM1 PE=3 SV=1 | 0.0487 | -0.1234 |
| F1MHP5 | Vascular endothelial growth factor receptor 3 OS=Bos taurus OX=9913 GN=FLT4 PE=3 SV=3 | 0.0398 | -0.1439 |
| A0A3Q1MPB2 | CCN family member 2 OS=Bos taurus OX=9913 GN=CCN2 PE=3 SV=1 | 0.0417 | -0.1488 |
| F1MZX6 | Myosin heavy chain 13 OS=Bos taurus OX=9913 GN=MYH13 PE=3 SV=3 | 0.0163 | -0.2357 |
| Q3ZCLO | Cysteine-rich secretory protein 2 OS=Bos taurus OX=9913 GN=CRISP3 PE=2 SV=1 | 0.0249 | -0.2901 |
| P68401 | Platelet-activating factor acetylhydrolase IB subunit beta OS=Bos taurus OX=9913 GN=PAFAH1B2 PE=1 SV=1 | 0.0090 | -0.3153 |
| Q3T114 | 2-iminobutanoate/2-iminopropanoate deaminase OS=Bos taurus OX=9913 GN=RIDA PE=2 SV=3 | 0.0269 | -0.3209 |
| A0A452DJ62 | Thrombospondin-4 OS=Bos taurus OX=9913 GN=THBS4 PE=3 SV=1 | 0.0499 | -0.3356 |
| A0A3Q1MB70 | Apolipoprotein M OS=Bos taurus OX=9913 GN=APOM PE=3 SV=1 | 0.0072 | -0.3472 |
| P00514 | cAMP-dependent protein kinase type I-alpha regulatory subunit OS=Bos taurus OX=9913 GN=PRKAR1A PE=1 SV=2 | 0.0493 | -0.3952 |
| A5D7V2 | PCDH12 protein OS=Bos taurus OX=9913 GN=PCDH12 PE=2 SV=1 | 0.0339 | -0.4057 |
| A0A3Q1MT60 | Macrophage migration inhibitory factor OS=Bos taurus OX=9913 GN=MIF PE=1 SV=1 | 0.0363 | -0.5381 |
| A0A3Q1M032 | Uncharacterized protein OS=Bos taurus OX=9913 PE=1 SV=1 | 0.0246 | -0.7632 |
| A5D9E8 | Mimecan OS=Bos taurus OX=9913 GN=OGN PE=2 SV=1 | 0.0452 | -0.8504 |

To determine if CAPG altered any of the *in vitro* ULF proteins, a comparison of proteins identified as differentially abundant in the conditioned medium compared to the unconditioned samples, and proteins identified in the CAPG treated samples as differentially abundant compared to vehicle controls samples was performed. Three proteins were identified as differentially abundant in both: ATL3 protein (ATL3), collagen type VI alpha 1 chain (COL6A1), and GST class-pi (GSTP1). ATL3 was increased (secreted) in the *in vitro* ULF, and further increased following exposure to rbCAPG (Table 3). COL6A1 and GSTP1 were both decreased in the conditioned medium compared to unconditioned, and COL6A1 was further decreased by rbCAPG, whereas GSTP1 was increased by rbCAPG.

**Table 3.** Proteins altered in conditioned medium and in response to rbCAPG. Proteins identified by TMT mass spectrophotometry in conditioned medium from vehicle control (VC) samples n=3 significantly differentially abundant (p<0.05) compared to unconditioned medium samples (n=2) or conditioned CAPG samples (rbCAPG added to medium, n=3) compared to conditioned VC samples. Conditioned medium obtained after flowing through a 3D microfluidic chip containing bEECs and bESCs. Medium collected after 24 hours from the bEEC compartment.

| Accession | Protein | Fold change VC vs unconditioned | Fold change CAPG vs VC |
| --- | --- | --- | --- |
| A5PJD6 | ATL3 protein (ATL3) | 3.275384538 | 0.5689 |
| E1BI98 | Collagen type VI alpha 1 chain (COL6A1) | -0.84089586 | -0.1099 |
| A0A3Q1LMV5 | GST class-pi (GSTP1) | -1.406021405 | 0.2374 |

### CAPG alters the endometrial transcriptome in vitro

Principal component analysis revealed that the largest contribution to transcriptional difference and clustering between samples was cell type (stromal or epithelial) as expected with epithelial cell samples clustering separately to stromal cells in PC1 (Figure 4B).

Exposure of epithelial cells to rCAPG under flow altered expression of 228 protein-coding genes (213 upregulated and 15 downregulated) and 6 long non-coding RNAs (5 upregulated and 1 downregulated) when compared to control (Supplementary Table S5). Similarly, rbCAPG addition altered 41 protein-coding genes in stromal cells when compared to the vehicle control samples (Supplementary Table S6). Twenty-three protein coding transcripts were commonly altered in both epithelial and stromal cell types (Figure 4C; Supplementary Table S7). All 23 shared transcripts were upregulated compared to vehicle control in both stromal and epithelial cell types.

The 228 protein coding genes altered by rbCAPG compared to vehicle control in epithelial cells in this 3D microfluidic system were subjected to biological processes gene ontology enrichment analysis (FDR <0.05), revealing that signal transduction in the absence of ligand, cell adhesion mediated by integrin, and muscle cell migration were the most highly enriched (Figure 4D: Supplementary Table S8). Enrichment analysis of the 41 protein-coding genes altered by the addition of rbCAPG when compared to vehicle control samples in stromal cells demonstrated that leukocyte migration, inflammatory response, and response to antibiotic were the most highly enriched biological process gene ontologies (Figure 4E: Supplementary Table S9).

### PDI alters the endometrium to support early pregnancy

The addition of rbPDI altered abundance of 18 proteins in the conditioned medium compared to vehicle control samples, with PDI the most highly abundant (Table 4). STRING DB analysis of 18 proteins revealed that many nodes had no connecting edges, and therefore no known interactions, but revealed some clear main interacting clusters, including a cluster comprised of DPP7, LGMN, and IFI30, another comprising LOC534578 and CD82, and another comprised of EIF5A and PGK1 (Figure 5A). Interestingly, PDI (termed P4HB in Figure 5A) was also shown to interact directly with two of the differentially abundant proteins secreted in response to PDI exposure-COL5A2 and CALU.

**Table4.** List of differentially abundant proteins in culture medium following PDI treatment. rbPDI added to the culture medium flowing through the fluidic bovine endometrium-on-a-chip device for 24 hours and proteins present in the conditioned medium determined by mass spectrophotometry, entially abundant proteins identified compared to vehicle control samples (n=3 biological replicates).

| Accession | Description | t-test | fold change |
| --- | --- | --- | --- |
| F6Q9Q9 | Protein disulfide-isomerase OS=Bos taurus OX=9913 GN=P4HB PE=1 SV=1 | 0.0015 | 2.0823 |
| Q2NL00 | Glutathione S-transferase theta-1 OS=Bos taurus OX=9913 GN=GSTT1 PE=2 SV=3 | 0.0250 | 1.6519 |
| A0A3Q1MDT9 | Collagen type V alpha 2 chain OS=Bos taurus OX=9913 GN=COL5A2 PE=4 SV=1 | 0.0462 | 1.5827 |
| A0A452DJJ3 | Glycerol-3-phosphate dehydrogenase [NAD(+)] OS=Bos taurus OX=9913 GN=GPD1 PE=3 SV=1 | 0.0012 | 1.3968 |
| A7MBB0 | VCAM1 protein OS=Bos taurus OX=9913 GN=VCAM1 PE=2 SV=1 | 0.0091 | 0.5930 |
| A0A3Q1NC75 | Calumenin OS=Bos taurus OX=9913 GN=CALU PE=1 SV=1 | 0.0180 | 0.3470 |
| F6QCC9 | Tetraspanin OS=Bos taurus OX=9913 GN=CD82 PE=1 SV=1 | 0.0112 | 0.3384 |
| E1BNR9 | Semaphorin 7A OS=Bos taurus OX=9913 GN=SEMA7A PE=3 SV=2 | 0.0452 | 0.2576 |
| A0A140T8D4 | Asparaginyl endopeptidase OS=Bos taurus OX=9913 GN=LGMN PE=3 SV=1 | 0.0215 | 0.1834 |
| A6QNX2 | DPP7 protein OS=Bos taurus OX=9913 GN=DPP7 PE=2 SV=1 | 0.0356 | 0.1391 |
| A0A3Q1MFC3 | Protein S100 OS=Bos taurus OX=9913 GN=S100A11 PE=1 SV=1 | 0.0007 | 0.1049 |
| Q6EWQ7 | Eukaryotic translation initiation factor 5A-1 OS=Bos taurus OX=9913 GN=EIF5A PE=2 SV=3 | 0.0404 | 0.0497 |
| A0A3Q1LZQ5 | Gamma-interferon-inducible lysosomal thiol reductase OS=Bos taurus OX=9913 GN=IFI30 PE=3 SV=1 | 0.0403 | -0.0924 |
| Q3T0P6 | Phosphoglycerate kinase 1 OS=Bos taurus OX=9913 GN=PGK1 PE=2 SV=3 | 0.0317 | -0.1903 |
| A0A3Q1M0F4 | BPTI/Kunitz inhibitor domain-containing protein OS=Bos taurus OX=9913 GN=LOC404103 PE=4 SV=1 | 0.0469 | -0.4046 |
| G5E573 | Vertnin OS=Bos taurus OX=9913 GN=VRTN PE=3 SV=1 | 0.0090 | -0.7103 |
| P35445 | Cartilage oligomeric matrix protein OS=Bos taurus OX=9913 GN=COMP PE=1 SV=2 | 0.0467 | -0.8299 |
| P80425 | Fatty acid-binding protein, liver OS=Bos taurus OX=9913 GN=FABP1 PE=1 SV=1 | 0.0137 | -1.5629 |

**Figure 5.**
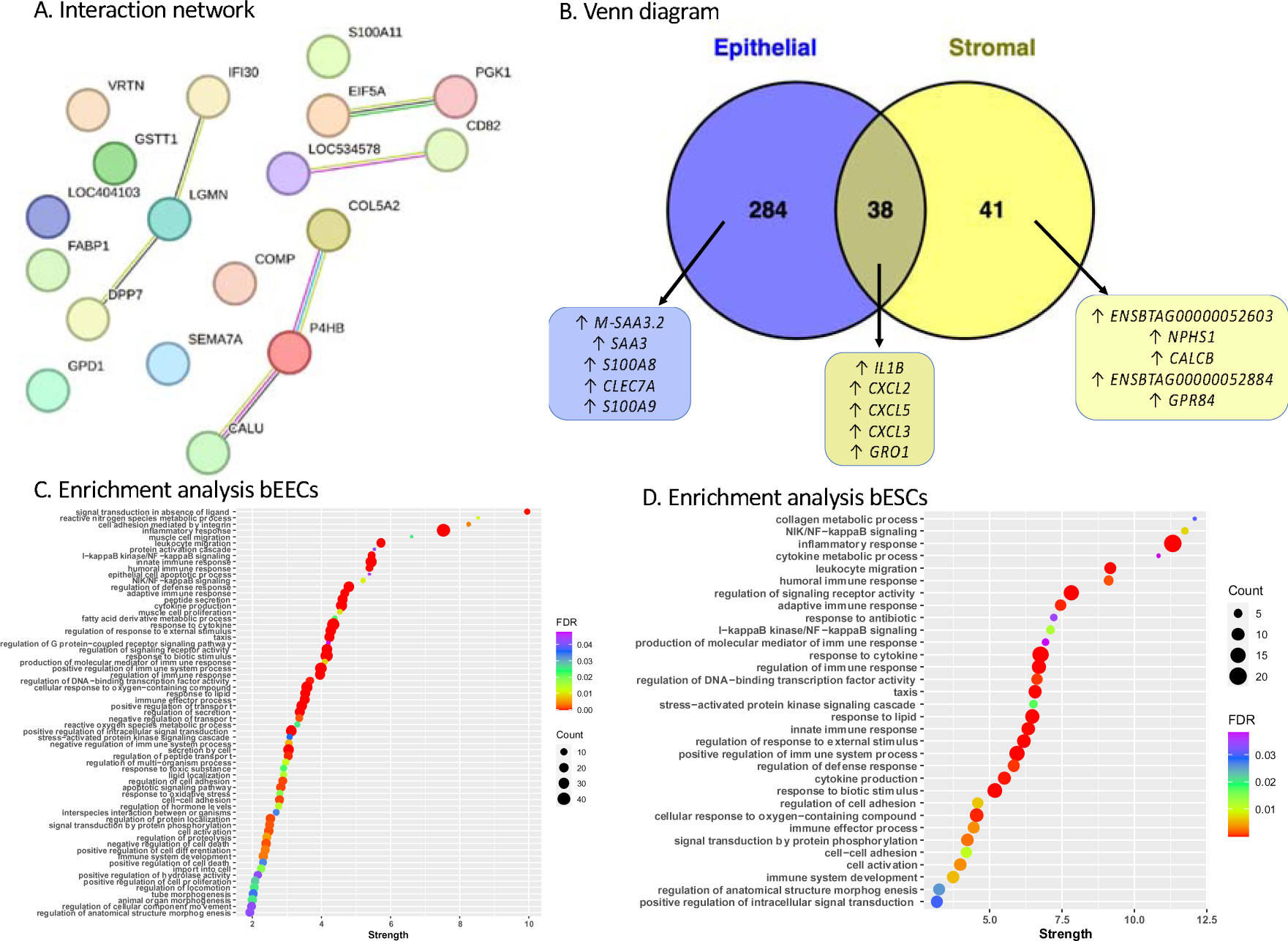
Impact of PDI on endometrial secretome and transcriptome. A. Interaction analysis of differentially abundant proteins in conditioned medium supplemented with rbPDI compared to vehicle control samples (p<0.05). Each node represents a protein, nodes with edges represent functional/physical protein associations with a minimum required interaction score of ‘medium 0.4’. PDI (P4HB in figure) was added to the culture medium. Thickness of edge represents the strength of supporting data. Produced using STRING DB. B. Venn diagram of significantly altered transcripts in response to rbPDI in different cell types within the bovine endometrium-on-a-chip microfluidic device. Significantly altered transcripts (padj<0.05, >1 or <-1 fold change) enrichment compared to vehicle control samples. ↑ upregulated, ↓ downregulated. Top 5 up/down regulated genes shown. Full data in supplementary table S12. C. Biological process gene ontologies enriched in bovine epithelial cells inside 3D microfluidic organ-on-a-chip device in response to rbPDI treatment. Analysis carried out in Webgestalt, FDR<0.05, strength indicates enrichment ratio. Full data in Supplementary table S13. Figure produced using ggplot in R Studio. D. Biological process gene ontologies enriched in bovine stromal cells inside 3D microfluidic organ-on-a-chip device in response to rbPDI treatment. Analysis carried out in Webgestalt, FDR<0.05, strength indicates enrichment ratio. Full data in Supplementary table S14. Figure produced using ggplot in R Studio.

To determine if PDI altered any of the *in vitro* ULF proteins, a comparison of proteins identified as differentially abundant in the conditioned medium compared to the unconditioned samples, and proteins identified in the PDI treated samples as differentially abundant compared to vehicle controls samples was performed. Three proteins were identified as differentially abundant in both analysis (Table 5): Calumenin (CALU), asparaginyl endopeptidase (LGMN), and fatty acid binding protein (FABP1). CALU was increased (secreted) in the *in vitro* ULF, and further increased following exposure to rbPDI. LGMN and FABP1 were both decreased in the conditioned medium compared to unconditioned, and FABP1 was further decreased by rbPDI, whereas LGMN was increased by rbPDI.

**Table 5.** Proteins altered in conditioned medium and in response to rbPDI. Proteins identified by TMT mass spectrophotometry in conditioned medium from vehicle control (VC) samples n=3 significantly differentially abundant (p<0.05) compared to unconditioned medium samples (n=2) or conditioned PDI samples (rbPDI added to medium, n=3) compared to conditioned VC samples. Conditioned medium obtained after flowing through a 3D microfluidic chip containing bEECs and bESCs. Medium collected after 24 hours from the bEEC compartment.

| Accession | Protein | Fold change VC vs unconditioned (in vitro ULF) | Fold change PDI vs VC |
| --- | --- | --- | --- |
| A0A3Q1NC75 | Calumenin (CALU) | 1.393152295 | 0.347 |
| A0A140T8D4 | Asparaginyl endopeptidase (LG MN) | -1.315570881 | 0.1834 |
| P80425 | Fatty acid-binding protein (FABP1) | -1.570223727 | -1.5629 |

### PDI alters the endometrial transcriptome in vitro

Exposure of epithelial cells to rbPDI altered expression of 314 protein-coding transcripts (295 increased and 19 decreased compared to vehicle control samples) and 8 long non-coding RNA transcripts (7 increased and 1 decreased compared to vehicle control samples) in the epithelial cells (Supplementary Table S10).

In stromal cells, the expression of 78 protein coding transcripts (75 increased and 3 decreased compared to vehicle controls) and 1 long non-coding RNA (increased in expression compared to vehicle controls) were significantly altered following rbPDI supplementation to the culture medium (Supplementary Table S11).

Venn diagram analysis demonstrated that 38 of the genes differentially expressed in response to rbPDI exposure were commonly altered in both cell types (Figure 5B: Supplementary Table 12). All commonly altered transcripts were upregulated compared to vehicle control in both epithelial and stromal cell types.

Enrichment analysis identified that the addition of rbPDI altered genes in the epithelial cells which were significantly enriched in biological processes when compared to vehicle control samples, such as signal transduction in absence of ligand, cell adhesion mediated by integrin, and inflammatory response (FDR <0.05, Figure 5C, Supplementary Table 13). Biological process GO terms enriched in the stromal cell compartment included collagen metabolic process, NIK/NF-kappaB signalling, and inflammatory response as the most highly enriched (FDR <0.05, Figure 5D, Supplementary Table 14).

To understand if there was a protein specific response in the different cell types we compared DEGs in response to CAPG and PDI exposed bEECs (Figure 6A) and bESCs (Figure 6B). 205 transcripts were commonly altered following rbCAPG and rbPDI treatment, whereas 29 transcripts were specifically in response to rbCAPG and 117 transcripts specific to rbPDI epithelial cells (Supplementary Table S15). In bESCs, 37 transcripts were commonly altered in response to rbCAPG and rbPDI treatment, whereas four transcripts were altered specifically in response to rbCAPG and 42 transcripts altered specifically in response to rbPDI (Supplementary Table S16).

**Figure 6.**
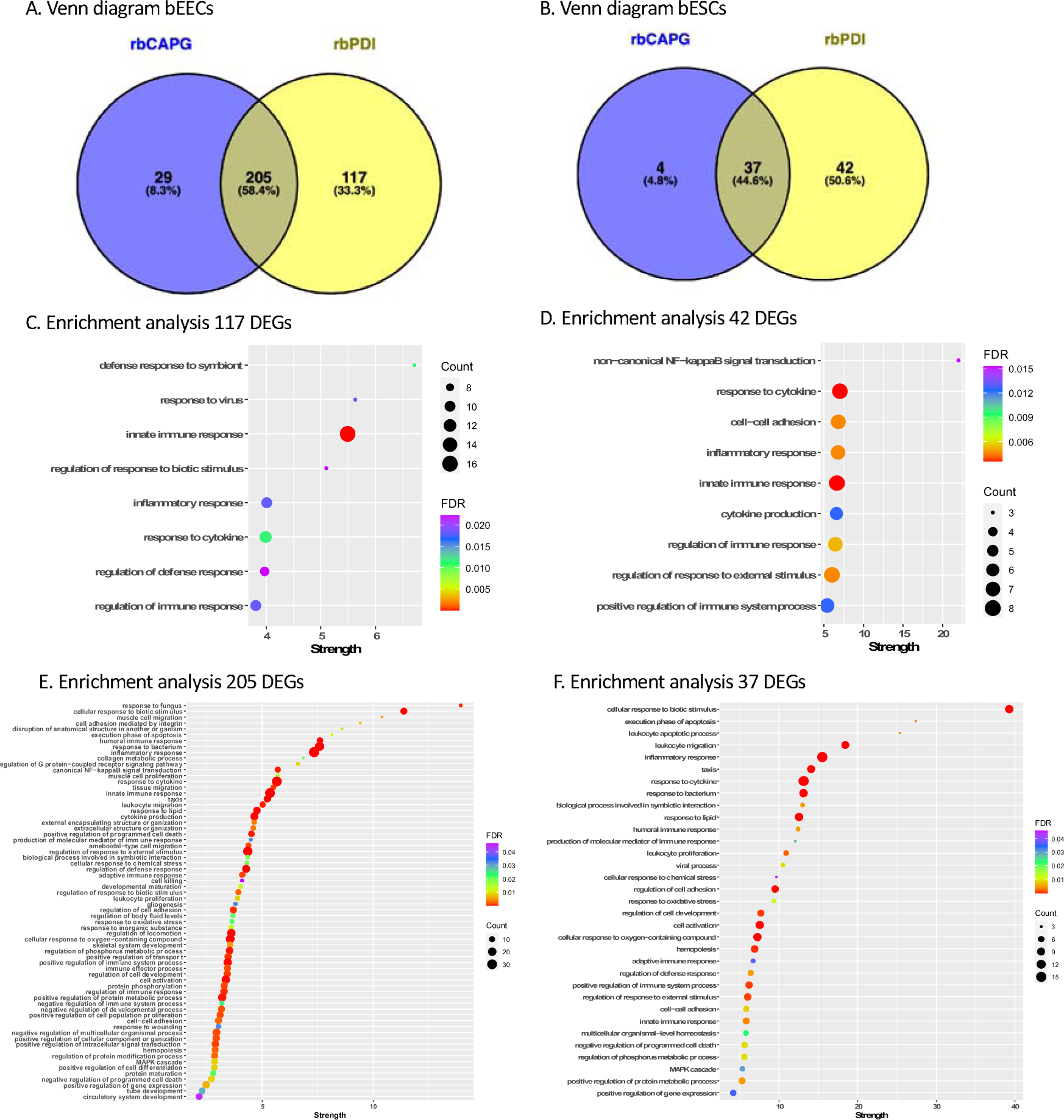
Response to CAPG and PDI is protein specific. Venn diagram comparison of rbCAPG and rbPDI induced DEGs in A) bEECs and B) bESCs. Cells treated with 1 μg/mL protein in 0.8 μL/min flow through the bEEC chamber of an endometrium-on-a-chip device also containing bESCs seeded on the other side of a porous membrane. DEGs determined in comparison to vehicle control samples, fold change >1 or <-1 and padj<0.05. Full data in supplementary table S15 and S16. Figures produced in Venny. C. Enriched gene ontology terms associated with 177 DEGs specific to rbDPI treatment but not rbCAPG treatment in bEECs. Cells treated with 1 μg/mL proteins under flow and DEGs determined in comparison to vehicle control samples, fold change >1 or <-1 and padj<0.05. GO enrichment analysis performed in webgestalt, non-redundant GO term dataset, FDR<0.05. Full data in Supplementary table S17. Figure produced in RStudio with ggplot2. D. Enriched GO terms associated with 42 DEGs specific to rbDPI treatment but not rbCAPG treatment in bESCs. Cells treated with 1 μg/mL proteins under flow and DEGS determined in comparison to vehicle control samples, fold change >1 or <-1 and padj<0.05. GO enrichment analysis performed in webgestalt, non-redundant GO term dataset, FDR<0.05. Full data in Supplementary table S18. Figure produced in RStudio with ggplot2. E. Enriched GO terms associated with 205 DEGs commonly altered by rbDPI and rbCAPG treatment in bEECs. Cells treated with 1 μg/mL proteins under flow and DEGS determined in comparison to vehicle control samples, fold change >1 or <-1 and padj<0.05. GO enrichment analysis performed in webgestalt, non-redundant GO term dataset, FDR<0.05. Full data in Supplementary table S19. Figure produced in RStudio with ggplot2. F. Enriched GO terms associated with 37 DEGs commonly altered by rbDPI and rbCAPG treatment in bESCs. Cells treated with 1 μg/mL proteins under flow and DEGS determined in comparison to vehicle control samples, fold change >1 or <-1 and padj<0.05. GO enrichment analysis performed in webgestalt, non-redundant GO term dataset, FDR<0.05. Full data in Supplementary table S20. Figure produced in RStudio with ggplot2.

Enrichment analysis on the 29 transcripts specific to rbCAPG in bEECs or the 4 transcripts specific to rbCAPG in bESCs did not reveal any significantly enriched GO terms. The 117 transcripts specific to rbDPI in bEECs revealed enrichment in eight GO terms, including defence response to symbiont, response to virus, and innate immune response (Figure 6C, Supplementary table S17). Enrichment analysis on the 42 transcripts specific to rbPDI treatment in bESCs revealed enrichment in nine GO terms, including non-canonical NF-kappaB signal transduction, response to cytokine, and cell-cell adhesion (Figure 6D, Supplementary table S18). Enrichment analysis on the 205 DEGs commonly elicited by both rbCAPG and rbPDI in bEECs identified 68 enriched GO terms, including response to fungus, cellular response to biotic stimulus, and muscle cell migration (Figure 6E, Supplementary table S19). Similarly, analysis on the 37 DEGs commonly altered by both rbCAPG and rbPDI in bESCs identified 33 enriched GO terms, including cellular response to biotic stimulus, execution phase of apoptosis, and leukocyte apoptotic process (Figure 6F, Supplementary table S20).

### Impact of culture model on endometrial response to conceptus-derived proteins

To understand how the 3D endometrium-on-a-chip culture system differs in responding to conceptus-derived factors, the transcriptional data from the static 2D cell cultures transcriptional response to rbCAPG (Tinning et al., 2020) and rbPDI (Tinning et al., 2024) were compared to data produced in this experiment. When epithelial cells were exposed to rbCAPG, 91 DEGs were commonly altered in both the 2D-static and 3D-flow systems, with 143 DEGs specific to the 3D-flow endometrium-on-a-chip system and 451 DEGs specific to 2D culture (Figure 7A: Supplementary Table S21). Secondly, the stromal cell transcriptional response from both systems were compared, revealing only 8 shared DEGs between both systems, with 33 DEGs specific to the 3D endometrium-on-a-chip system and 32 DEGs specific to the static culture system in response to rbCAPG (Figure 7B: Supplementary Table S22).

**Figure 7.**
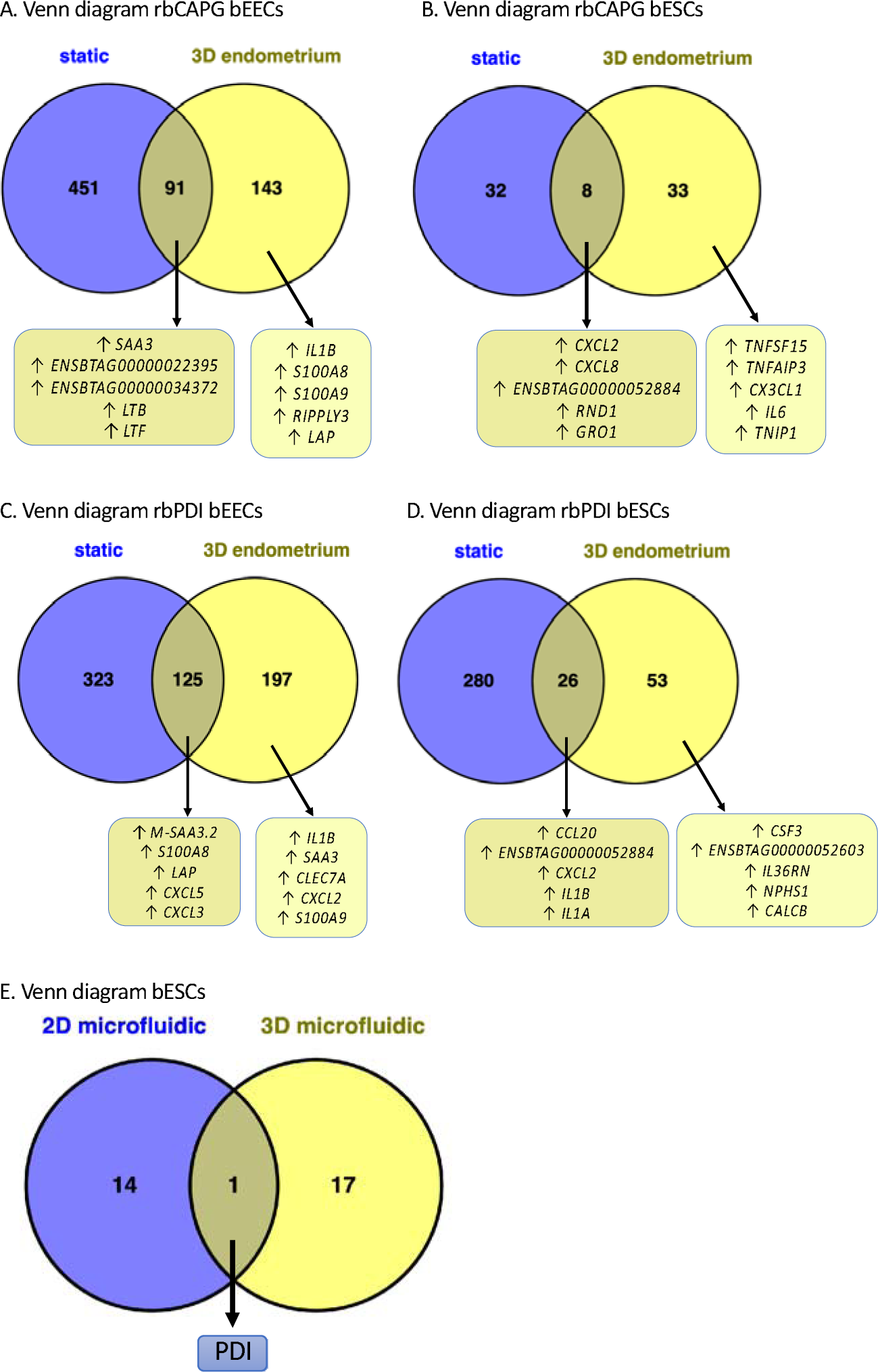
Comparison of differentially expressed genes or altered secretome from 2D-static vs 3D-endometrium-on-a-chip systems treated with rbCAPG and PDI. Static culture system was 2D and a single monolayer of cells presented in Chapter 2. 3D endometrium-on-a-chip system is a 3D culture of both epithelial and stromal cells, with rbPDI or rbCAPG applied to the epithelial cell side under flow 0.8 μL/min. Differentially expressed genes (padj <0.05, fold change >1 or <-1) determined following rbCAPG treatment in A. bEECs or B. bESCs compared to vehicle control samples, or following rbPDI treatment in C. bEECs or D. bESCs compared to vehicle control samples. ↑ upregulated, ↓ downregulated. Top 5 up/down regulated genes shown. Full data in Supplementary tables S21-S24. E. Comparison of the conditioned medium and secretome of 2D and 3D microfluidic culture systems following rbPDI treatment. 2D culture system from Tinning et al (2024) consisted of bovine endometrial epithelial cells in a simple microfluidic channel in a monolayer. The 3D microfluidic culture system presented here contained both stromal and epithelial cells applied to either side of a porous membrane. Proteins were identified by TMT mass spectrophotometry compared to vehicle control samples (p<0.05). Full data in Supplementary table S25.

In epithelial cells exposed to rbPDI, 125 DEGs were commonly altered between 2D-static and 3D-Flow systems, with a further 197 DEGs specific to the 3D-flow and 323 DEGs specific to 2D-static (Figure 7C: Supplementary Table S23). Secondly, the stromal cell transcriptional response from both systems were compared, revealing only 26 shared DEGs between both systems, with 53 DEGs specific to the 3D endometrium-on-a-chip system and 280 DEGs specific to 2D static culture (Figure 7D: Supplementary Table S24). Finally, the secretome of the cells cultured within the 3D endometrium on a chip in response to rbPDI (Table 5) was compared to the secretome of epithelial cells cultured in a 2D microfluidic system in response to rbPDI (Tinning et al., 2024). No commonly altered proteins were identified, 17 proteins were specifically altered in the 3D endometrium on a chip system, and 14 proteins specifically altered in static 2D culture (Figure 7E: Supplementary table S25).

## DISCUSSION

Classic 2D static culture monocultures do not allow us to investigate the role of conceptus-derived proteins upon the endometrium in way that recapitulates the *in vivo* endometrial environment. To understand how the conceptus-derived proteins may alter the transcriptome and secretome of the endometrium *in vivo,* a 3D endometrium on a chip system was developed to co-culture bEECs and bESCs and expose the bEECs to conceptus-derived factors under flow. A 3D bovine multicellular endometrium-on-a-chip *in vitro* device was utilised in a microfluidic system to model the bovine endometrial tissue to aid in deciphering the complex cross-talk between endometrium and conceptus. Firstly the ‘*in vitro*’ ULF secretome was identified and similarities to the *in vivo* ULF proteins identified (Forde et al., 2015), and secondly the 3D endometrium-on-a-chip transcriptional and secretory response to conceptus-derived factors (CAPG and PDI) was investigated and shown to be different to that demonstrated using 2D *in vitro* culture systems.

### The in vitro uterine luminal fluid

The 3D endometrial model presented here is comprised of two chambers separated by a porous membrane allowing cellular communication through the membrane via secreted factors such as extracellular vesicles and secretory proteins/miRNAs etc. The design of this endometrium-on-a-chip is that the static chamber represents the underlying stromal compartment, and the microfluidic channel under flow represents the uterine lumen and epithelial monolayer lining the endometrial tissue. This design was used to recapitulate the communication that occurs between conceptus and endometrium *in vivo.* The bovine conceptus elongates to fill the length of the ipsilateral horn by day 16, both horns by day 21 (Betteridge et al., 1980; Kastelic et al., 1988). The outermost trophectoderm cells of the conceptus would be closely aligned with the luminal epithelium during this period of pregnancy recognition and peri-implantation. Therefore, the luminal epithelial cells would be exposed to any conceptus-derived secretions first. Due to the orientation of cells in this endometrium on a chip system, the conceptus derived factors could be applied under gentle flow, bathing the epithelial cells. This allowed the chip to be used here to investigate the endometrial response to conceptus-derived factors in a manner which mimics the endometrial tissue structure and exposure to conceptus-derived factors. Collection of the conditioned medium also gave an insight into the secretome of the epithelial cells within the system (representing the ULF *in vitro*).

### ULF proteins secreted by the 3D in vitro endometrium are involved in adhesion and supporting endometrial development

One of the key components of this study was to understand how this novel 3D endometrium on a chip system compared to static and *in vivo* produced ULF and transcriptional responses. Enrichment analysis of the endometrial secreted proteins into the epithelial-facing system ‘lumen’ revealed many GO terms associated with cell-to-cell adhesion, cell-to-matrix adhesion, cell-substrate adhesion, cell migration, tissue development, and structure development. This indicates that many of the proteins secreted into the *in vitro* ULF may be involved in the establishment and development of the endometrial tissue within the system. The endometrium took around 3-4 days to develop to the point of 90% confluence of the epithelium, and then was left intact for another 24 hours. In terms of tissue establishment this is relatively short, and likely resulted in tissue-like compartments which were still primarily undergoing self-organisation, forming an intact monolayer of epithelial cells (forming cell-cell adhesions), and attaching to the underlying substrate (the membrane and/or ibiTreat Ibidi polymer material) (Xavier da Silveira dos Santos and Liberali, 2019). Other studies attempting to recapitulate the 3D bovine endometrium *in vitro* which have assessed tissue structure used culture lengths of 14 days (MacKintosh et al., 2015) and 35 days (Díez et al., 2023). Further culture of the cells prior to using the system may have resulted in a more mature tissue and a different secretome, although it was visualised that a longer period of culture led to a change of the morphology of the stromal cells seeded in the upper static chamber. Establishing the ideal length of culture time and/or optimising the culture medium may produce a secrete which better matches that of the *in vivo* produced ULF.

### In vitro bovine ULF has limited similarity to in vivo ULF

Proteomic analysis of the spent conditioned culture medium flowed through the microfluidic channel revealed that 68 proteins were actively secreted by the *in vitro* endometrium-on-a-chip system on the epithelial side (mimicking the ULF secretion). When compared to a published dataset of day 16 non-pregnant bovine ULF proteome (Forde et al., 2015), only 5 proteins were shared with the 68 identified here. Approximately half of the proteins identified in the study by Forde *et al* were successfully converted to the same protein identifier used here, meaning it is possible that a greater number of proteins were shared with the proteins secreted in this *in vitro* system but were lost in the data processing steps.

The first of the 5 identified proteins both *in vitro* in the 3D system and in *in vivo* ULF, keratin 19, has previously been identified as being increased in abundance in the ULF from day 8 pregnant cattle compared to cyclic cattle (Muñoz et al., 2012), and of higher expression in bovine embryos (*in vitro* fertilised) compared to lower competency nuclear transfer embryos (Pfister-Genskow et al., 2005). Keratin 19 may therefore be secreted to support conceptus development in cases of lower competency.

Annexin A2 is involved in attachment of murine blastocysts *in vitro,* and infusion of annexin A2 artificially to the murine uterus increased the number of implantation sites (B. Wang et al., 2015). Annexin A2 has also shown to be upregulated during the oestrus phase (cycle day 20-22 post ovulation) compared to the diestrus phase (cycle day 5-15 post ovulation) in cattle, which the authors attribute to rising P4 concentration in the maternal circulation acting upon the endometrium (Bauersachs, 2005). Conversely in humans, annexin A2 was found to be more highly expressed in the endometrium during the midsecretory phase (approximately cycle day 21 post menses start, and approximately day 7 post ovulation) compared to the late proliferative phase (cycle day 8-10 post menses start) (Kao et al., 2002). The midsecretory phase corresponds to the stage at which the human embryo implants (day 6-9 post ovulation). This may point to an important role in the endometrial production of annexin A2 in supporting implantation, although at different stages of the cycle and embryo development in different species, which is retained in the 3D *in vitro* endometrial model presented here. A future possibility could be applying hormones to the system to replicate the oestrus cycle to investigate the impact on the transcriptome and/or secretome.

CD9 is a common exosome membrane marker, and has been found in the proteome of exosomes from the bovine ULF in multiple studies (Qiao et al., 2018; Piibor et al., 2023). CD9 being found in the *in vitro* ULF from the 3D endometrium-on-a-chip system indicates that the endometrium-on-a-chip is functioning to secrete exosomes/EVs as the endometrium does *in vivo,* and that many of the proteins identified here may be packaged within EVs.

The protein basal cell adhesion molecule was secreted in this *in vitro* system and is an immunoglobulin adhesion molecule-involved in cell-cell adhesion (Wai Wong et al., 2012). It was also found highly differentially expressed in the endometrium compared to the conceptus in ovine, linked to interact with secretory protein LAMA5 (Yang et al., 2022). Although little is currently known about any potential roles of basal cell adhesion molecule in the context of the endometrium or embryo, it is viable that it is involved in cell-cell adhesion of the conceptus and endometrium during implantation, or in maintaining/remodelling the endometrium during the implantation period.

Finally, pre-mRNA-processing factor 6 enhances the activation of the androgen receptor and is also a component of the spliceosome which functions to splice precursor mRNAs to form mature mRNAs within the cell (Liu et al., 2021). Androgen receptor signalling has been associated with endometrium development and uterine gland formation in mice knockouts (Choi et al., 2015) and placental function (Parsons and Bouma, 2021). Pre-mRNA-processing factor 6 may be secreted into the ULF to support either spliceosome function or androgen receptor activation in the conceptus, endometrium, or placenta.

Although the 3D endometrial model presented here doesn’t recapitulate the *in vivo* secreted ULF, it does retain some candidate proteins which are also secreted *in vivo* which may be of importance and constitutively secreted during the peri-implantation period. The composition of ULF varies throughout the oestrus cycle (Mullen et al., 2012; Faulkner et al., 2013; Mullen et al., 2014; Simintiras et al., 2019) and by pregnancy stage (Forde, Simintiras, et al., 2014; Forde, McGettigan, et al., 2014; Sponchiado et al., 2019), therefore the ULF is variable dependent upon endometrial exposure to maternal hormones and the influence of the conceptus. The 3D endometrium on a chip system presented here could be adapted to investigate the influence of steroid hormone exposure (by adding the upper stromal ‘maternal-side’ chamber) or conceptus derived factors (by adding to the lower epithelial ‘conceptus-side’ channel. The device can also be used in reverse, with the ‘maternal-side’ stromal cells in the lower flow channel to recapitulate the maternal circulatory system, and has been used to investigate the influence of maternal glucose and insulin concentrations (De Bem et al., 2021), which further influence the secretome in the system. To summarise, this model could be optimised by accounting for other factors which influence the endometrial secretome *in vivo* and can be used to investigate the effect of different factors upon the endometrium secretome *in vitro.* The model was further used here to investigate the role of the conceptus-derived factors, CAPG and PDI, in modulating the endometrium.

### CAPG alters the secretome and proteome of the 3D in vitro endometrium-on-a-chip to support early pregnancy processes

When rbCAPG was added to the microfluidic flow through on the endometrial epithelial-side (replicating the uterine luminal side) it was to replicate how CAPG would be secreted from the conceptus and how the endometrium would be exposed to CAPG *in vivo.* This resulted in a change in the secretome of the *in vitro* ULF produced of which they interacted with each other, around GAPDH as a central connecting node in the network. GAPDH has been identified in bovine non-pregnant uterine EVs and ULF (Piibor et al., 2023), and also in pregnant ULF on day 16 and in greater abundance on day 19 in cattle (Forde, McGettigan, et al., 2014). CAPG may therefore act upon the endometrium to stimulate the production of GAPDH to trigger a network of other proteins being secreted.

One of those proteins, MIF (macrophage inhibitory factor), was previously identified in the bovine (Forde et al., 2015) pregnant ULF as a ‘non-classical ISG protein’ (i.e. not associated with a type I interferon response), secreted by bovine epithelial cells *in vitro* in response to IFNT (Wang and Goff, 2003), and shown to be secreted by human endometrial cells in response to hCG (Akoum et al., 2005). MIF’s main function is as a pro-inflammatory cytokine, and has been discussed to have potential roles in many areas of reproductive biology (Jovanović Krivokuća et al., 2021). Interestingly, MIF has been shown to stimulate migration and invasion of trophoblast cell lines *in vitro* (Jovanović Krivokuća et al., 2015), as well as many roles in supporting placental development and function (Jovanović Krivokuća et al., 2021). Therefore, CAPG may be secreted by the conceptus to stimulate the endometrium to secrete MIF, alongside IFNT (or hCG in humans), to support implantation and placentation.

Three other proteins which were secreted in response to CAPG which were shown to interact-OGN, THBS4, and COL6A1. OGN is a growth factor expressed only in the endometrium and not in the bovine conceptus, THBS4 expressed as a ligand on the bovine conceptus, and COL6A1 a ligand expressed in both the bovine conceptus and endometrium (Mamo et al., 2012). CAPG may activate secretion of these proteins to further promote physical interactions between the conceptus and the endometrium and to stimulate conceptus growth during elongation.

A comparison of the proteins differentially abundant *in vitro* within the 3D endometrium on a chip system (*in vitro* ULF proteins), to those altered following exposure to rbCAPG revealed three overlapping proteins-ATL3 was increased in abundance in the *in vitro* ULF and also following rbCAPG exposure. ATL3 is a receptor GTPase involved in the endoplasmic reticulum promoting tubular fusion and promotes degradation of the tubular endoplasmic reticulum (Chen et al., 2019). ATL3 has not yet been discussed in the literature the context of uterine luminal fluid. The other two proteins (COL6A1 and GSTP1) were decreased in the *in vitro* ULF, indicating that the cells were taking up the proteins from the culture medium rather than secreting them as ULF proteins.

### CAPG may alter the endometrial transcriptome to mediate the immune response and remodel the endometrium

The transcriptome of the endometrial epithelial and stromal cells was altered in response to rbCAPG when compared to the vehicle controls samples. One the most highly positively differentially expressed genes in epithelial cells in response to rbCAPG was *MMP13*, a matrix metalloproteinase, which is involved in degrading collagenous extracellular matrix. *MMP13* expression has been shown to be modulated at the site of implantation in pigs (Yoo et al., 2023), and extracellular matrix remodelling is highly involved in implantation and placentome formation in cattle (Yamade et al., 2002). *CCL2* was also differently positively expressed in response to rbCAPG specifically in epithelial cells. *CCL2* has been shown to be more highly expressed in pregnant endometrium in cattle than non-pregnant on days 15 and 18 of pregnancy, but the authors found that this wasn’t in response to IFNT in an *ex vivo* explant culture (Sakumoto et al., 2017). *CXCL8* was specifically more highly expressed only in stromal cells in response to rbCAPG, and has also been found to be increased in response to IFNT (Sakumoto et al., 2017). Other C-X-C motif chemokines, including *CXCL2*, *CXCL3*, and *CXCL5,* were commonly more highly expressed in both cell types in response to rbCAPG. Chemokines have been demonstrated to regulate endometrial-conceptus interactions in cattle (Sakumoto et al., 2017), pigs (Złotkowska and Andronowska, 2019), and humans (Zhang et al., 2022). *TNFSF15* was specifically upregulated in stromal cells only, but has previously been identified as a conceptus-specific cytokine (Mamo et al., 2012).

In the endometrial epithelial cells, the biological processes most enriched amongst genes differentially expressed in response to rbCAPG compared to vehicle control samples were mostly surrounding immune- and cytokine-related processes, but also cell adhesion, muscle cell migration/proliferation, and secretion. Cell adhesion is a process required for the attachment of the conceptus trophectoderm to the endometrial epithelium (implantation) (D’Occhio et al., 2020). Muscle cell proliferation and migration may be related to the epithelial-stromal cell communication to promote endometrial remodelling during early pregnancy. Finally, secretion of factors from the endometrium epithelium is essential for histotrophic nutrition available in the ULF for the developing conceptus. CAPG may promote secretion from the endometrial epithelium to enrich the ULF.

Overall, CAPG may be secreted by the conceptus to support immune regulation, secretion of ULF, epithelial-stromal cell communication, and implantation processes in cattle.

### PDI exposure may modify the endometrial transcriptome to mediate the inflammatory response to conceptus

Exposure to rbPDI altered the transcriptome of both epithelial and stromal cells in the bovine 3D endometrium-on-a-chip system, eliciting a greater number of DEGs specifically in epithelial cells. Some of the most highly expressed transcripts in each cell type are involved in processes such as immune response and regulation. The 5 most highly expressed genes in epithelial cells in response to rbPDI (*M-SAA3.2, SAA3, S100A8, CLEC7A, S100A9)* are all linked to immune response (Oliveira et al., 2010; Swangchan-Uthai et al., 2013; Murata et al., 2020). PDI also induced C-X-C motif chemokine expression in both cell types, similarly to rbCAPG. Specifically, in stromal cells the expression of *CALCB*, involved in placental development, was increased in response to rbPDI. *CALCB* has been identified as expressed during the ‘window of receptivity’ in human endometrium (Dorostghoal et al., 2017).

Many biological processes were enriched amongst the genes differentially expressed in response to rbPDI, including many involved in immune response, secretion, and adhesion in epithelial cells. Similarly immune response and adhesion related biological processes were enriched in stromal cells. Despite a smaller number of DEGs being altered in response to rbPDI in stromal cells than epithelial cells, the biological processes enriched are very similar, except for those related to secretion. This is likely due to epithelial cells being responsible for secretion of the histotrophic contribution to ULF *in vivo,* which nourishes the pre-implantation conceptus. PDI therefore may modulate the immune response to conceptus *in vivo and* modify the secretome to support conceptus development.

### PDI also modifies the endometrial secretome in vitro

Supplementation of the culture medium with rbPDI altered the secretome of the 3D endometrium-on-a-chip system. Two proteins found to be secreted in response to rbPDI were also shown to directly interact with PDI by interaction analysis-COL5A2 and CALU. COL5A2 has been identified as an endometrial receptivity gene upregulated in response to the embryo in human (Haouzi et al., 2011) and during implantation in rabbit (Liu et al., 2016). COL5A2 is a fibrillar type of collagen, associated with ECM organisation, therefore PDI may promote COL5A2 secretion to act in an autocrine manner upon the endometrium to support implantation.

PGK1 was secreted in response to rbPDI and has previously been found to be secreted *in vitro* in response to human trophoblast cell line-derived EVs (Muhandiram et al., 2023), and is thought to be involved in decidualisation of human stromal cells (Tong et al., 2018). PGK1 could support endometrial remodelling during the implantation period in cattle. Another protein, LGMN has previously been identified as being upregulated in the bovine endometrium during early pregnancy and speculated to be involved in placentome formation as it is a protease activator and could therefore be involved in endometrial remodelling (Ledgard et al., 2009).

A comparison of the proteins differentially abundant *in vitro* within the 3D endometrium on a chip system, to those altered following exposure to rbPDI revealed three overlapping proteins-CALU was increased in abundance in the *in vitro* ULF and also following rbPDI exposure. Extracellular CALU reduced matrix metalloproteinase-13 cleavage of fibulin-1 ((Q. Wang et al., 2015). Fibulin-1 is an extracellular matrix protein which is more highly abundant in human endometrial glands during the proliferative phase but more highly abundant in the stroma during the secretory phase, indicating it may have a role in implantation (Nakamoto et al., 2005). The other two proteins (LGMN and FABP1) were decreased in the *in vitro* ULF, indicating that the cells were taking up the proteins from the culture medium rather than secreting them as ULF proteins.

### PDI and CAPG differentially altered the transcriptome in the endometrium on a chip system

A comparison of the transcriptomic response of the epithelial and stromal cells cultured within the 3D endometrium on a chip system revealed that many (45-58% of total DEGs) of the differentially expressed genes were commonly altered between both rbCAPG and rbPDI exposure. This indicates that the proteins may have overlapping functions *in vivo*. rbCAPG altered minimal (29 transcripts in bEECs and 4 transcripts in bESCs) DEGs which weren’t also altered by rbDPI. Comparing previously published data also demonstrates an overlap in the transcriptional response to rbCAPG (Tinning et al., 2020) and rbPDI (Tinning et al., 2024). Enrichment analysis did not reveal any enriched GO terms among those limited DEGs. However, rbPDI altered more transcripts which were not altered by rbCAPG (117 in bEECs and 42 in bESCs). Enrichment analysis on those 117 DEGs revealed many enriched GO terms, all of which were related to immune system regulation in bEECs. The 42 DEGs specific to rbPDI identified GO terms including cell-cell adhesion and non-canonical NF-kappaB signal transduction. Non-canonical NF-kappaB signalling is associated with negative regulation of type I interferons (Jin et al., 2014). Therefore, PDI may have a role in modulating IFNT production *in vivo*.

### 3D endometrium-on-a-chip system demonstrates both differences and similarities in secretome and transcriptome response to conceptus-derived factors

Of the differentially expressed genes elicited by CAPG/PDI in the 3D endometrium on a chip system, between 61-80% (varied between cell types and treatment) of those were specific to the 3D culture system, whereas 20-39% of transcripts were also altered in the static 2D systems (Tinning et al., 2020; Tinning et al., 2024). This indicates that the endometrial response to conceptus-derived factors differs when under flow and when epithelial cells are exposed to stroll cells (recapitulating the in vivo endometrial structure), compared to under standard static culture techniques. This reinforces our need to develop 3D in vitro models to study endometrial-conceptus communication to better model the *in vivo* environment. One gene found highly expressed in response to rbCAPG in the 3D culture system only was *RIPPLY3*, which has previously been found to be upregulated in the endometrium during the window of implantation in pregnant mice (Aikawa et al., 2022).

The secretome of the 3D endometrium-on-a-chip from the epithelial side was wholly different from that of the 2D epithelial-only channel in response to rbPDI (Tinning et al., 2024). This demonstrates that the 3D co-culture of stromal and endometrial cells changes how they function, as evidenced by work showing stromal cells supports epithelial cell growth and differentiation in human *in vitro* cultures (Arnold et al., 2001), further supporting our need for in vitro culture systems which consider and recapitulate aspects of the in vivo environment.

In conclusion we have demonstrated that a 3D bovine microfluidic endometrium-on-a-chip was successfully utilised to mimic the endometrium, ULF secretion, and exposure to conceptus-derived factors. The *in vitro* ULF produced from the endometrial side of the 3D bovine endometrium-on-a-chip system had limited similarity to *in vivo* day 16 non-pregnant ULF, but exposure to conceptus-derived factors, PDI and CAPG, altered the secretome and transcriptome of the bovine endometrium-on-a-chip in a protein-specific manner. The endometrial response to rbPDI was also shown to differ in the 3D system compared to a previously used 2D microfluidic systems, indicating the importance of using co-culture systems.

## ACKNOWLEDGEMENTS

Research in NF’s lab is supported by N8 agri-food pump priming, QR GCRF, as well as BBSRC grant numbers BB/R017522/1, BB/X007332/1 and Wellcome Trust grant 227178/Z/23/Z.

## DATA AVAILABILITY

All transcriptomic data is currently being uploaded to GEO (GEO xxx). All proteomic data will be available on Proteome Exchange (XXX).

